# Large-scale genome-wide association study of food liking reveals genetic determinants and genetic correlations with distinct neurophysiological traits

**DOI:** 10.1101/2021.07.28.454120

**Authors:** Sebastian May-Wilson, Nana Matoba, Kaitlin Wade, Jouke-Jan Hottenga, Maria Pina Concas, Massimo Mangino, Eryk J. Grzeszkowiak, Cristina Menni, Paolo Gasparini, Nicholas J. Timpson, Maria G. Veldhuizen, Eco de Geus, James F. Wilson, Nicola Pirastu

**Affiliations:** Centre for Global Health Research, Usher Institute, University of Edinburgh; Department of Genetics, University of North Carolina at Chapel Hill; UNC Neuroscience Center, University of North Carolina at Chapel Hill; Population Health Sciences, Bristol Medical School, University of Bristol; Medical Research Council (MRC) Integrative Epidemiology Unit (IEU) at the University of Bristol; Dept of Biologial Psychology, FGB, Vrije Universiteit Amsterdam; Amsterdam Public Health research instiute, Amsterdam UMC; Institute for Maternal and Child Health—IRCCS, Burlo Garofolo; Department of Twin Research and Genetic Epidemiology, King’s College London; NIHR Biomedical Research Centre at Guy’s and St Thomas’ Foundation Trust; Department of Medicine, Surgery and Health Sciences, University of Trieste; Department of Anatomy, Faculty of Medicine, Mersin University; Amsterdam Public Health research insitute, Amsterdam UMC; MRC Human Genetics Unit, Institute of Genetics and Molecular Medicine, University of Edinburgh

## Abstract

Variable preferences for different foods are among the main determinants of their intake and are influenced by many factors, including genetics. Despite considerable twins’ heritability, studies aimed at uncovering food-liking genetics have focused mostly on taste receptors. Here, we present the first results of a large-scale genome-wide association study of food liking conducted on 161,625 participants from UK Biobank. Liking was assessed over 139 specific foods using a 9-point hedonic scale. After performing GWAS, we used genetic correlations coupled with structural equation modelling to create a multi-level hierarchical map of food liking. We identified three main dimensions: high caloric foods defined as “Highly palatable”, strong-tasting foods ranging from alcohol to pungent vegetables, defined as “Learned” and finally “Low caloric” foods such as fruit and vegetables. The “Highly palatable” dimension was genetically uncorrelated from the other two, suggesting that two independent processes underlie liking high reward foods and the Learned/Low caloric ones. Genetic correlation analysis with the corresponding food consumption traits revealed a high correlation, while liking showed twice the heritability compared to consumption. For example, fresh fruit liking and consumption showed a genetic correlation of 0.7 with heritabilities of 0.1 and 0.05, respectively. GWAS analysis identified 1401 significant food-liking associations located in 173 genomic loci, with only 11 near taste or olfactory receptors. Genetic correlation with morphological and functional brain data (33,224 UKB participants) uncovers associations of the three food-liking dimensions with non-overlapping, distinct brain areas and networks, suggestive of separate neural mechanisms underlying the liking dimensions. In conclusion, we created a comprehensive and data-driven map of the genetic determinants and associated neurophysiological factors of food liking beyond taste receptor genes.

## Introduction

Food consumption is one of the most important factors influencing our health and contributes to a large amount of excess mortality in the world ^1^. With the near limitless availability of food in the Western world due to mass distribution, there has been a shift in factors driving dietary behaviour from merely consuming the food that is available to one of choice. For this reason, in parallel to understanding the effect of food consumption on health, there has been an increasing interest in understanding the drivers behind people’s choices in order to direct them toward being more nutritious and thus reduce the burden of various diseases. Food choice is a complex process which involves many different factors such as personal preferences, health status, ethical beliefs and context. Rather than measures of preference (or choice), liking of foods reflects the individual hedonic response to foods^2^ and is closely related to biology^3–5^. Thus, understanding food liking may be the first critical step in designing better, more targeted dietary interventions and more acceptable nutritious foods.

Food liking is a complex trait clearly influenced by biology, psychology^6^, the surrounding environment^7^, branding^8^, culture^9^ and genetic inheritance^10^. In particular, twin studies have shown that food preferences are moderately heritable traits, with around 50% of their variance in children being explained by genetic factors plus mostly shared environmental effects^11,12^. In adults, while heritability remains stable, the shared environmental component disappears in favour of the non-shared one (eg personal experiences)^13–16^

Although several recent GWAS have looked at the genetic variants associated with food consumption^17–19^, when it comes to liking attempts to identify the genetic factors underlying these food-liking traits have focused mostly on candidate gene studies^20^ (PMID: 22888812) (e.g., genes encoding taste receptors such as TAS2R43 and coffee liking^21^), with mixed results^22^. More recently, genome-wide approaches have been used to identify several genes related to the liking of different foods in an untargeted manner. For example, genetic variants have now been identified as being associated with the liking of sweet foods^23^ or more common foods^24^ such as cilantro/coriander^25^. However, these studies have focused either on specific sensations/tastes or tend to be small in sample size and are so underpowered to detect the likely modest effect sizes of common genetic variation on more specific food-liking traits.

Here, we present the results of a genome-wide association study (GWAS) for detailed food- and beverage-liking traits in more than 150,000 participants from the UK Biobank study, with replication in up to 26,154 individuals across 11 independent cohorts. Furthermore, we used genetic correlations combined with genomic structural equation modelling to create a multi-level map of the relationships between different food preferences, highlighting three main domains that we define as “Highly palatable”, “low caloric” and “learned” foods. We show that these dimensions are genetically correlated to distinct brain areas, behavioural, socio-economic, anthropometric, and biochemical traits which are expected to correlate with these food-liking factors, indirectly validating the model. Finally, we unravel the pleiotropic effects of many of the identified genetic variants, mapping them to the food-liking traits they influence directly.

## Methods

### Study populations

#### UK Biobank

Analyses were conducted on data collected in the UK Biobank study under project 19655. UK Biobank recruited more than 500,000 people aged 37 to 73 years from the United Kingdom between 2006 2010. The study, participants, and quality control have been described previously^26^. All subjects gave written informed consent. UK Biobank was approved by the North West Multi-Centre Research Ethics Committee (MREC) and in Scotland, UK Biobank was approved by the Community Health Index Advisory Group (CHIAG). We included only subjects who completed the food liking questionnaire and were of European descent. Full details of the genetic information, food-liking phenotypes are presented below. Genotyping was conducted using the UK Biobank or the UK BiLEVE Axiom Arrays. (Affymetrix, Santa Clara, CA, USA). Further details about imputation, principal components analysis, and QC procedures can be found elsewhere (https://biobank.ctsu.ox.ac.uk/crystal/crystal/docs/genotyping_qc.pdf).

### Food-liking phenotypes

Food-liking traits were collected through an online questionnaire comprising 152 items, including both foods and non-food items, which was administered in 2019 to all UK Biobank participants who had agreed to be recontacted by the study. The questionnaire is an extension of the one previously used in Pallister et al. 2015^13^ and Vink et al. 2020^15^. Given that the questionnaire was administered online to participants pictures were removed, and we used a 9-point Hedonic scale^27^, where 1 corresponds to “Extremely dislike” and 9 to “Extremely like”. Other options also included “Have never tried it” and “Prefer not to answer”. Details of the questionnaire can be found at (https://biobank.ndph.ox.ac.uk/showcase/showcase/docs/foodpref.pdf). Of the 152 items, only the 139 pertaining to food and drink were retained for this specific study, while those which referred to habits such as physical activity were not included. Coffee- and tea-liking were measured asking both with and without sugar, we thus defined two additional measures for each: the first was the maximum score given to coffee and tea (coffee max and tea max) to reflect liking for the drink in the preferred way; the other was instead estimated as the difference between the sweetened vs the unsweetened drink to reflect polarization in liking, so higher values meant a higher liking for the sweetened drink while negative numbers reflected a stronger liking for the unsweetened drink.

A full list of the food-liking traits used in the study, mean number of participants and standard deviation of responses can be found in Supplementary Table 1.

### Statistical analyses

#### GWAS

Genome-wide association analysis was performed for each of the 144 food-liking traits using the raw reported score rescaled so that values would range between 0 and 1. After regressing each food liking trait with age, sex and the first 10 genetic principal components, array type and batch, we accounted for genetic relatedness between the participants using GRAMMAR+ residuals^28^ as estimated in fastGWA^29^. Finally, GWAS was performed using regscan^30^ assuming an additive model on all SNPs with MAF > 0.001. Given the high number of food-liking traits analysed and the high correlation between them, to estimate study-wide significance, we first estimated the minimum number of independent components which accounted for at least 95% of the variance over all the traits. This was achieved by estimating the eigen decomposition of the genetic correlation matrix between all the studied food liking questionnaire items. We estimated that 34 components are sufficient to explain >95% of genetic variance and we thus considered a p-value of p<1.47 x 10^-9^ (5 x 10^-8^ / 34) as the study-wide significant threshold. Given that many loci showed association with multiple traits, we also considered all associations that reached a conventional genome-wide significance threshold (p<5×10^-8^) if the SNPs were in the same genomic locus as a study-wide significant one.

### Clustering of food-liking items, hierarchical model construction

To describe the interrelationships between the food-liking questionnaire items we used hierarchical factor analysis where multiple steps of factor analysis are performed. In our case we first estimated the pairwise genetic correlations between all pairs of the original food-liking items from the questionnaire using the LD-score regression (ldsc) software^31^. We then performed hierarchical clustering using Ward’s D2 method, as implemented in the hclust function of R. We then visually defined a first set of groups that showed a high level of within-group correlation across the individual food-liking items. We then estimated a first set of factors, one for each defined group of items. Validity of each of these models were estimated using GenomicSEM R package^32^ and looking at goodness of fit metrics, specifically comparative fit index (CFI) >0.9 and a Standardized Root Mean Square Residual (SRMR) <0.1. If the model did not have a good fit, we checked whether this could be due to single items and they were removed accordingly. Once the first level of factors was defined, we estimated the effect of each SNP on the factor variable, obtaining for each factor complete GWAS summary statistics. We then estimated genetic correlations between the resulting factors, if any two factors exhibited a genetic correlation larger than 0.9, the items of the two groups were merged together and a new overall factor was estimated. The factors GWAS then become the starting point for building a higher order of factors. This procedure was repeated until we ended up with a hierarchical structure composed of only 4 high order factors and up to 4 levels. To make the results more readable we assigned to each of the factors a label to better interpret what it is capturing (e.g. Meat for the factor derived from all the meat items), however to keep the difference between observed and derived factor traits, we have added an “F” before the label (e.g. F-Meat)

### Estimation of the effect of each SNP with each factor

To estimate the effect of each SNP on each of the latent variables or factors, we first used GemonicSEM to estimate the loadings of each observed variable onto the latent one. We then applied the method described in Tsepilov et al 2020^33^. Briefly the effect of each SNP on each factor is estimated as the weighted linear combination of the effect of the SNP on each index variable, where the weights are represented by the loadings of each item on the latent variable. This is analogous to using the usergwas function in GenomicSEM, but leads to a large reduction in computing time.

### Comparisons between food liking and food consumption traits

In order to understand how our food-liking measures were related to diet, we performed genetic correlation analysis between the GWAS of the food frequency questionnaire and the alcohol consumption data, available through the Pan UKBB project website (https://pan.ukbb.broadinstitute.org/). We also compared heritability (h^2^) estimated using LD-score regression. Heritability comparison and genetic correlations analysis was limited to those traits for which either the exact same item was present in both the food frequency questionnaire and the food liking questionnaire (e.g. white wine) or items with a corresponding and similar item between both questionnaires (e.g. Cheese).

### Genetic correlations with other complex traits

Genetic correlations with other complex traits for the three top order traits was performed using the ldhub web portal (http://ldsc.broadinstitute.org/ldhub/). Given the high number of correlations estimated, we selected a set of 31 traits representative of the socio-economic, anthropometric, blood biochemistry and health-related behaviour traits, to summarise the results.

### Locus definition and colocalisation analysis

To define the boundaries of each locus, we first selected all SNPs with p-value <1×10^-5^ and then estimated the distance between each consecutive SNP located on the same chromosome. Two consecutive SNPs were identified as belonging to different loci if they were more than 250 kb apart. This approach allows locus identification based on peak shape rather than a fixed distance from a sentinel SNP. A locus was then considered “significant” if it contained at least one SNP with p-value below 1.47×10^-9^. Loci which showed overlapping boundaries were merged. To finally test if the underlying causal SNPs between the merged loci were the same or were just close to each other in the genome, we utilised the HyPrColoc method^34^. Briefly, HyPrColoc tests if a group of traits (e.g., food liking traits) colocalise and returns the probability of each SNP in the locus being causal. Moreover, it returns a separate overall regional colocalisation probability. We thus divided the positional loci into sub-loci based on the results of this analysis and then used the SNP with the highest probability of being causal for each cluster as sentinel SNP.

### Meta-analysis and replication

Replication of the GWAS for the questionnaire items was conducted using up to 26,154 samples coming from 11 different cohorts mostly of European ancestries: ALSPAC, INGI-CARL, INGI-VB, INGI-FVG, CROATIA-Korcula, NTR, Silk Road, the TWINS UK cohort, CROATIA-Vis and VIKING. Details of each cohort can be found in Supplementary Table 2

Given that each cohort used a related but different questionnaire meta-analysis was performed only on the overlapping food liking traits for which at least 10,000 samples were available.

Given that different cohorts have used different scales we have rescaled the results so that they would reflect a scale going from 0 to 1. Prior meta-analysis QC on the summary stats was performed using EasyQC v 28.3^35^.

All traits were meta-analysed using inverse variance weighting conducted using METAL v 2018-08-28^36^.

Given that only a limited number of traits was available for at least ten thousand samples it was possible to attempt replication of only 235 SNP-trait associations.

### Gene prioritisation

To define the gene that was most likely to be responsible for the observed association at each locus, we proceeded with custom prioritisation according to the following criteria. We first ran haploR v.4.0.2^37^ using r^2^=0.8 as the threshold using the sentinel SNP in each sub-locus. If a SNP was not available within the HaploReg resource, we used the most likely available one. Then, genes were prioritised if the locus met one of the following conditions (in order of importance):

1. The sentinel SNP is itself or is in strong LD (r^2^>0.8) with a non-synonymous SNP in the gene;
2. The sentinel SNP is itself or is in strong LD (r^2^>0.8) with a coding SNP in the gene (synonymous or in the untranslated region of the gene);
3. The top SNP is intronic or is in complete LD with an intronic SNP in the gene;
4. The top SNP is in strong LD (r^2^>0.8) with an intronic SNP in the gene;
5. The closest gene.

### Estimating the direct effect of each SNP on specific food-liking and latent factor traits

One of the aims of this study was to understand which SNPs influence different food-liking traits and if these associations were mediated through some higher order latent factor or if it was directly influencing the food trait of interest. For example, if we consider alcoholic beverages, we can imagine that some SNPs may influence liking of lower order food traits such as beer or wine through overall liking of alcohol, or directly on beer- liking or both. We thus aimed at untangling the direct effect of the SNPs on each food-liking and latent factor trait, from those mediated through other connected traits.

To do this, we used GenomicSEM, which allows fitting the effect of each SNP onto multiple traits at the same time, while considering their relationships. The limitation, however, is that it is not possible to fit the effect of the SNP on all observed variables and the latent variable at the same time, given that the number of observed SNP estimates is less than the parameters we need to estimate.

Therefore, we developed a strategy that enabled us to get all the required estimates. To illustrate this strategy, let’s imagine we have 3 correlated food-liking traits (T1-T3), for which a SNP effect is available and where the common variance can be explained by a latent variable L1 (Fig 1 Panel (A- 1)). The first step of our analysis was to estimate the effect of the SNP on the latent variable L1 (Fig 1 Panel (A-2)); to fit the effect of the SNP on all 4 traits at once to estimate all 4 parameters, we need to provide at least the same number of observed estimates. However, only 3 are available. To solve this, we created a new model, where we considered L1 as an observed variable and created a new dummy latent variable (DV) that explained all 4 traits and that was highly correlated (0.99) with L1. The SNP effect is then fit onto the original 3 food-liking traits (T1-T3) and the dummy variable such that we could obtain the estimate of the SNP effect on the latent variable and the residuals of the 3 food-liking traits at the same time.

**Fig. 1.**
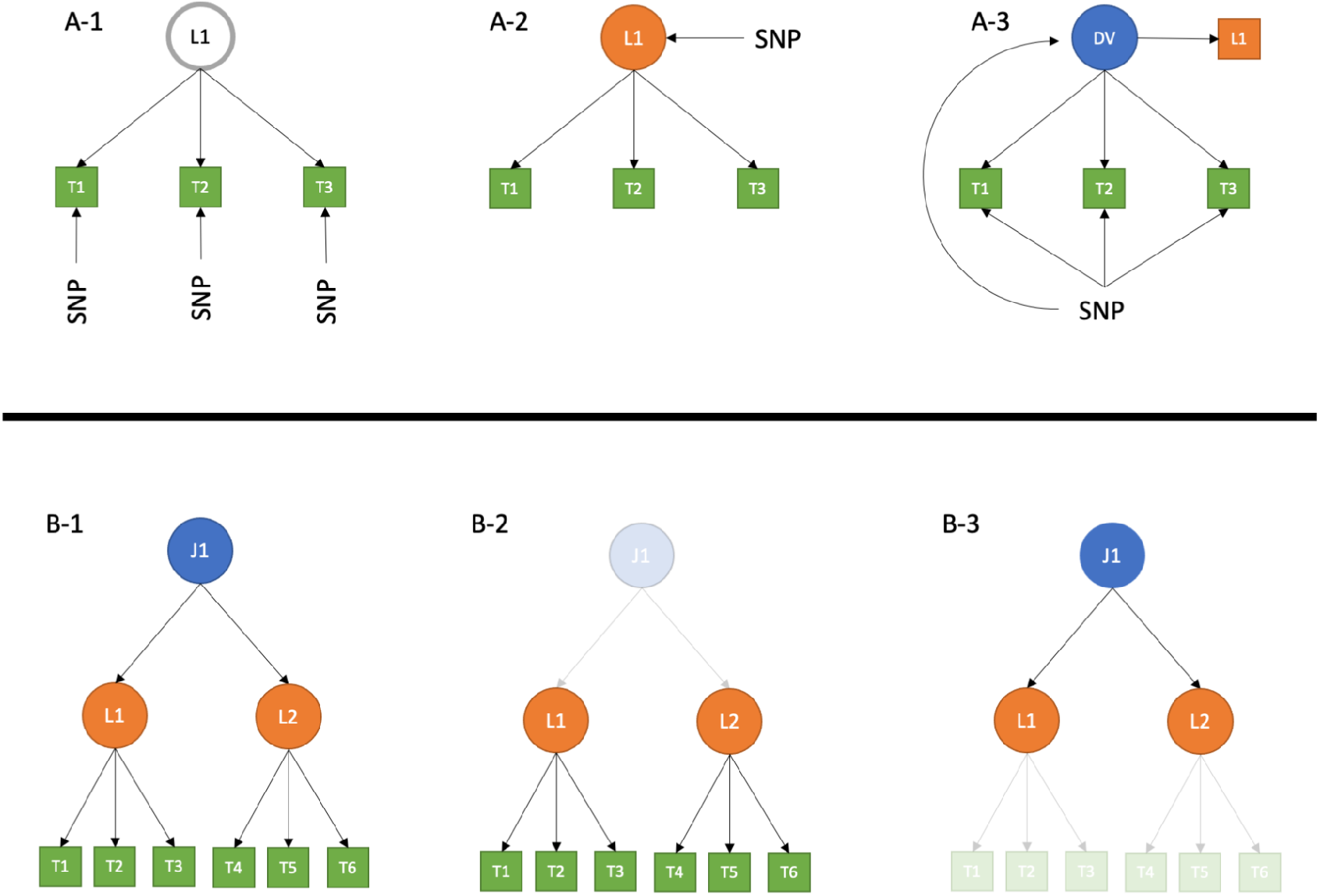
Strategy to map loci to specific traits. Panel A shows the strategy to fit the SNP effect contemporarily on all food liking traits in the model. We started with the SNP effect on each observed trait participating in the model (A-1). We then used GenomicSEM to estimate the effect of the SNP on the latent variable, L1, based on the observed ones (A-2). We finally used the SNP estimate on L1 as though it were directly observed and created a new dummy latent variable (DV) strongly correlated to L1 (0.99) and fit the SNP effect on LD and all participating food liking traits at the same time (A3). Panel B shows the strategy used to fit the multiorder model. The full model (B-1) is split into levels composed of 1 latent variable and its observable variables and the strategy described in panel A is applied. This is repeated level-by-level (B-3) and then results of all conditioning models for each trait are compared.

The described approach is useful to solve simple one factor models, but it cannot be directly applied to the complex hierarchical model we created, as it would be computationally infeasible. We thus split the hierarchical model of food items into smaller trees, where only one latent variable and its observable food traits were used. In efforts to retain the overall structure, we fixed the loadings of the food-liking traits onto the factor to be the same as those estimated during the construction of the model. Fig. 1 panel B summarises this strategy.

For all intermediate order traits, this approach led us to have for several factors 2 different conditional estimates: one where the latent factor trait was conditioned on the index food traits and another in which it represented the index trait. To select which estimate captured best the direct effect, we select the one with the smallest absolute value of Z-score. We can imagine that if the effect of the SNP is mediated through another trait, conditioning on this trait will lead to a decrease in the effect, and thus the estimate with the smallest effect would correspond to the correct one. Fig. 1 panel B1-3 reports a scheme of this strategy. To test if the conditional SNP estimate was different from the original estimate we used the method from Clogg et al 1995^38^:

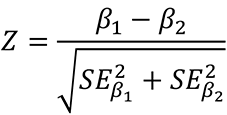

We considered “direct effect only” SNP/trait effects which showed p>0.05 at this test.

### Functional and Tissue enrichment analysis

For enrichment analysis we expanded the gene selection to all those which were mapped to loci which were associated with at least one of the food liking traits at p<5×10-8. Information about the full list of loci can be found in Supplementary Table 3.

Tissue enrichment analysis was conducted using FUMA^39^ looking at the general and specific GTEx tissues as reference. Gene Ontology term enrichment analysis was conducted using the enrichGO() function from the clusterprofiler R package (3.16.1)^40^.

### Correlation with brain MRI traits

To estimate genetic correlation with brain MRI, we first obtained 3,260 GWAS summary statistics on Imaging-derived phenotypes (IDP) from multimodal brain imaging (excluded diffusion MRI and ICA25) from Oxford Brain Imaging Genetics Server - BIG40 (https://open.win.ox.ac.uk/ukbiobank/big40/)^41^. These IDPs included morphological traits as well as functional neural response traits. For the morphology measures cortical thickness, surface area and volumes were calculated in regional brain areas for various parcellations of the brain (Freesurfer atlases).

Briefly these areas/networks were derived by applying a technique called “group independent component analysis” (ICA) which identifies a prespecified number of networks as independent from each other as possible. This was estimated in UK Biobank using two different values: 25 and 100 with the ICA100 identifying smaller brain areas. In particular for our analyses we used the ICA100 traits which include 55 non-artifact nodes and 1485 edges (between nodes) for a total of 1540 traits.

The functional neural response traits included the average neural response over time during a resting-state scan in 55 non-artifact network maps from the ICA100 IDPs (each encompassing multiple regional brain areas), as well as the edges between all 55 ICA maps. The derivation of the ICA100 traits has been described in detail elsewhere^42^. We removed IDPs with low heritability or large uncertainty of heritability estimates (p < 0.05), resulting in 2,329 IDPs tested for genetic correlations. Genetic correlations were estimated using high-definition likelihood (HDL)^43^ to maximise power. Genetic correlations were tested only with the three main dimensions coming from the hierarchical factor analysis. We applied FDR to correct multiple testing on 6987 pairs (significance threshold was set to q<0.05) (Supplementary Table 4).

## Results

Supplementary Table 1 presents descriptive summary statistics for the food-liking traits.

### Mapping the relationships between food items

As the first step in our analysis, we aimed to map the relationships between the different food preferences. After running the GWAS on all the questionnaire items, we computed the genetic correlation matrix and compared it with the phenotypic one (Fig S1). The resemblance between the two correlations was very high (r=0.91, Supp Fig 1B), but the genetic correlations between the food-liking traits were on average twice as large as the phenotypic correlations, likely due to the high measurement error in the food-liking questionnaire.

Looking at the hierarchical clustering of the foods based on their genetic correlations (Supplementary Fig 1A), two main groups of foods were easily identified: one that included what could be considered “high-reward” foods, such as meat, desserts and fried foods, and another group that included a larger and wider variety of items ranging from fruit, to alcoholic beverages, unsweetened caffeinated drinks and cheese.

Hierarchical factor analysis as described above led to a tree structure model composed of up to 4 levels (Fig 2A and supplementary file 1), with three main dimensions of food liking at the top with the final model comprising 119 questionnaire items out of the initial 144.

**Fig 2.**
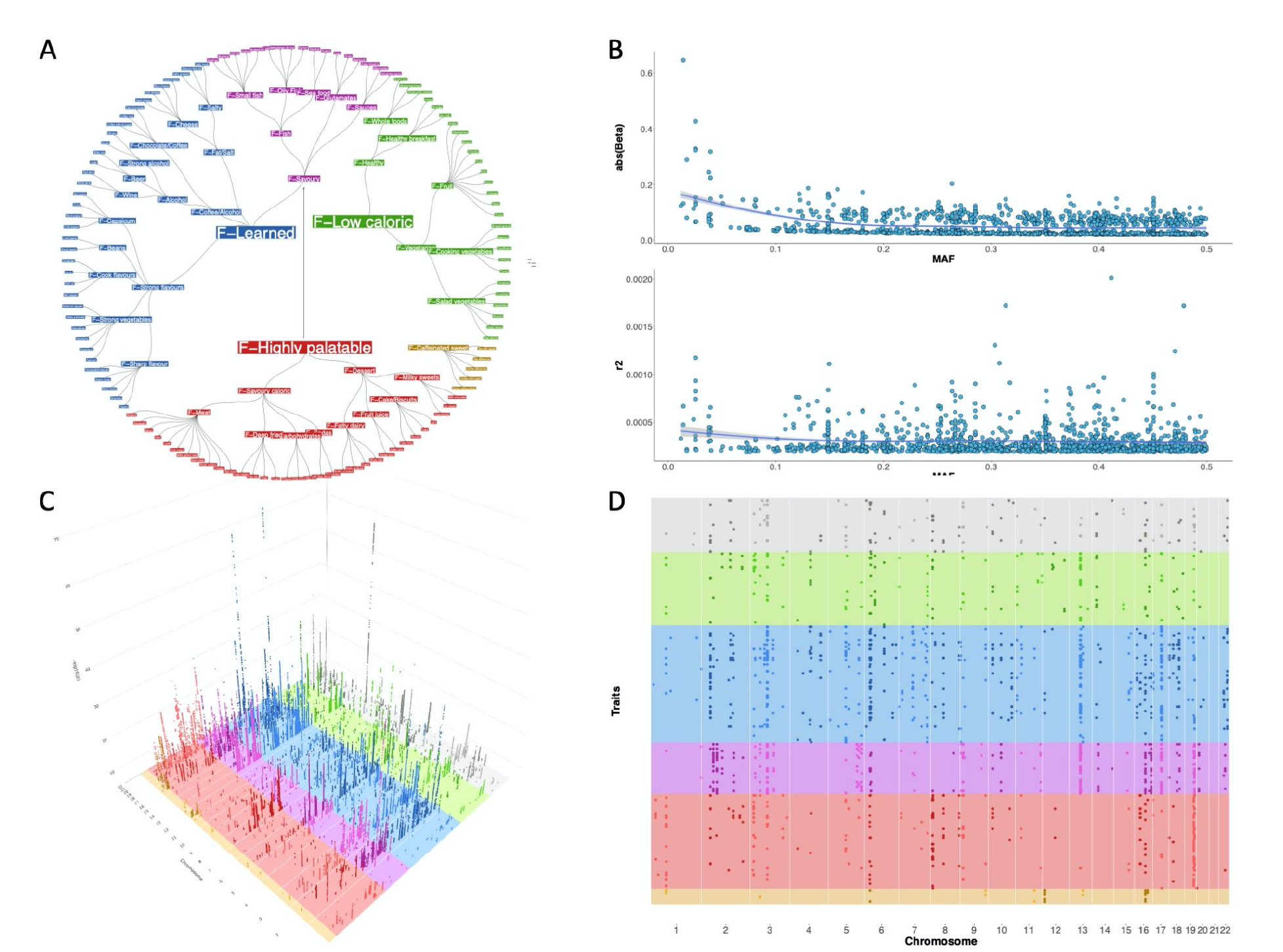
Food-liking map and genome-wide association results. (A). Hierarchical model of relationships between liking of different foods. The leaves represent the original food liking traits which were measured with the questionnaire. Colours reflect the membership in one of the four independent dimensions: Red, F-Highly palatable; Blue, F-Learned; Green, F-Low Caloric; Light brown, F-Caffeinated sweet drinks. F-Savoury foods are colored purple as they contribute to both F-Highly palatable and F- Learned Foods. (B). Upper panel represents the relationship between the minor allele frequency and effect size. As in most complex traits, there is an inverse relationship between MAF and effect size. Lower panel represents the same SNPs but r^2^ is reported on the y axis, showing no relationship between the two measures. (C). 3D Manhattan plot, only SNPs with p<5×10^-8^ have been reported. Colours reflect those used in panel A. (D). Bird’s-eye view of the Manhattan plot. Each dot represents the top SNP from each of the sub-loci.

The first factor trait included highly energetically rewarding and widely accepted foods such as desserts, meat and savoury foods which we named “F-Highly palatable”. The second was composed mainly of low caloric foods such as vegetables, fruit and wholegrain, which we defined as “F-Low caloric”. The third was composed of items for which liking is generally acquired, such as unsweetened coffee, alcohol, cheese and strong-tasting vegetables, which we refer to as “F-Learned”. Finally a fourth minor group was composed of F-sweetened caffeinated drinks.

F-Low caloric and F-Learned traits showed a moderately strong genetic correlation (rG = 0.59), while the F-Highly palatable trait was more or less completely independent from either (rG, 0.05 and 0.16, respectively). Finally the F-Caffeinated Sweet Drinks showed a weak positive correlation with the F-Highly palatable dimension (rG =0.39) and a weak negative correlation with the F-Learned and F-Low caloric groups (rG=-0.3 and rG= -0.25, respectively).

### Genetic Correlation with food consumption

Overall, we detected a very strong correlation between the liking measures and their corresponding consumption traits (Fig 3, Supplementary Table 5), with all correlation coefficients being >0.7, with the exception of beer (rG=0.4) and white bread (rG=0.1). Looking at heritability estimates, the mean SNP heritability for the liking traits (∼0.08) was double that for the consumption traits (∼0.04), and food liking always showed higher values, with the exception of dried fruit, where there was little evidence of a difference and tea, where heritability was higher for consumption.

**Fig 3.**
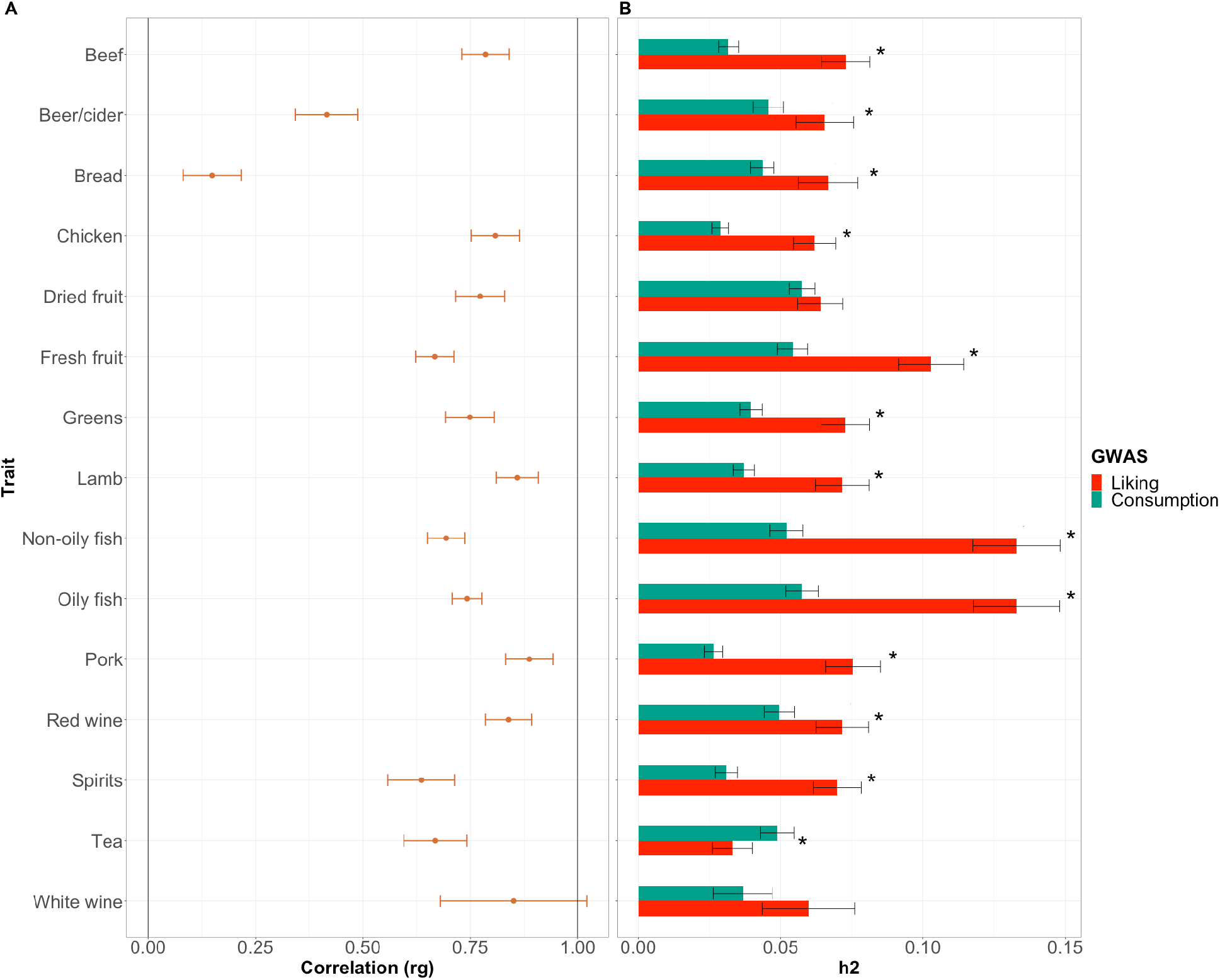
Genetic comparison between food liking and food consumption traits. Panel A. reports the genetic correlations between consumption and liking of the same food for all foods for which both were available, bars represent 95% CI. Panel B. Comparison between SNP heritability of food consumption (red) and liking (green). Bonferroni-corrected significant differences are indicated with a star.

### Genetic correlation with other complex traits

Genetic correlations with other complex traits (Fig 4 and Supplementary File 2) showed differences between the three main F-traits. The F-highly palatable trait showed correlations with higher indices of obesity (higher BMI and body fat percentage), lower socioeconomic status and lower levels of physical activity despite showing a positive correlation with non-sedentary jobs. F-Highly palatable was also correlated with higher sodium and creatinine in urine, likely reflective of a diet richer in protein and added salt. The F-Low caloric trait showed positive correlation with higher physical activity and use of dietary supplements but also with a non-sedentary job suggesting that people reporting higher liking for the F-Low caloric trait show a general tendency for a “healthier” lifestyle. This is reflected also by the negative correlation with urinary sodium and creatinine suggestive of a healthier diet and with lower body fat percentage. The F-Learned trait was positively correlated with indexes of higher socioeconomic status such as years in schooling and a sedentary job, a overall healthier blood lipid and obesity profile and higher physical activity although it also correlated with higher likelihood of having smoked and higher alcohol consumption.

**Fig 4:**
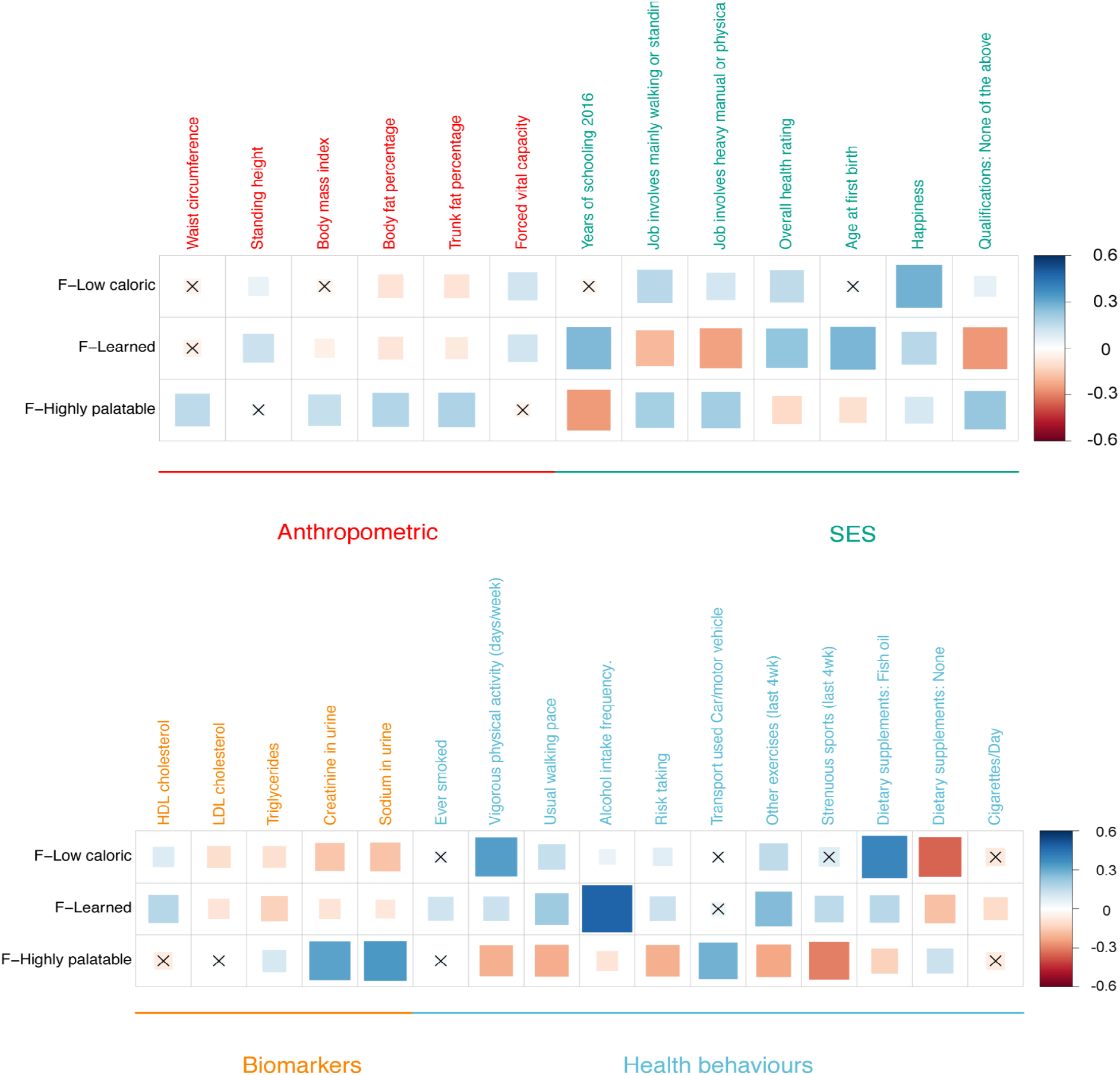
Genetic correlation between the three main food liking factors and other selected complex traits. X indicates FDR > 0.05.

### GWAS results

In our GWAS of food liking, we identified evidence for 1401 genetic associations divided into 173 loci (Fig 2, Supplementary Table 6). 143 loci out of 173 corresponding to 1270 out of 1401 associations showed correlations with multiple traits, with the *FTO* locus being associated with 58 traits, suggesting high levels of pleiotropy.

### Pleiotropy and colocalisation

Colocalisation analysis with HyperColoc (Supplementary Tables 7 and 8) showed that most traits that were associated in the same locus, also colocalised. Within the 143 loci, 138 showed at least one group of traits which colocalised with each other for a total of 203 distinct clusters. 225 of the 1270 association did not colocalise with any other trait.

### Replication

Replication analysis in up to 26,154 people (median 15,736) from 11 different cohorts was able to replicate 61 (one tailed p<0.05 and same direction of effect) out of 235 testable associations (26%) (Supplementary Table 9). However, 194 associations corresponding to 82.5% showed consistency of direction of effect (binomial test p=5×10^-25^).

### Gene prioritization

Gene prioritization (see Methods for details) allowed us to identify 250 genes as being most likely causal. Close to half of the associations (43.8%) were intragenic, with roughly 7% of non-synonymous variants and about the same proportion (∼6%) of SNPs located either in the 3’ or 5’ untranslated region. Only ∼1% could be explained by synonymous variants. Rather unsurprisingly, 12 of the prioritised genes encoded either taste (4) or olfactory receptors (8) and highlighted many novel associations. For example, the strongest association we detected was between *OR4K17* and liking of onions (beta=0.31 on a 9 point scale, p=4 x 10^-71^).

Amongst taste receptors, associations were identified only for bitter receptors and all were associated to traits belonging either to the learned or low caloric group while none were associated with the Highly palatable foods. A similar pattern was observed also for the genes encoding olfactory receptors. Of particular interest are the variants of the *TAS2R38* gene, which were associated with salty foods, alcoholic beverages, horseradish and grapefruit, confirming our previous results^17,44^, which provided evidence for association between this locus and adding salt to food and consuming red wine, but also expanding this finding to other alcoholic beverages.

Similarly, there were other cases which corroborated and expanded upon previous reports. For example, variants near the *FGF21* gene, which has been previously associated with consumption of sweet foods^45^, were also negatively associated with stronger-tasting foods, especially fish but also eggs, mayonnaise and fatty foods.

### Distinguishing direct from mediated effects

As shown by the colocalization analysis, the hierarchical relationships between the food preference traits give rise to a very high level of pleiotropy. Thus, in order to be able to predict the potential function of the identified genes, it is important to be able to understand at which level of the hierarchical tree of food liking the variant is primarily associated with. If we think of liking fruit, for example, we can imagine that some variants may be associated with all fruits while others may be associated with specific fruits such as apples or oranges. To resolve this issue, we fit the effect of each sentinel SNP onto all nodes of the model at the same time as outlined in Materials and Methods and determine if the observed effect was direct or mediated through one of the correlated traits. Of the initial 1261 associations which could be tested within the hierarchical model, only 495 were inferred to be direct effects. As an exemplar case, Fig 5 shows the effects of this approach for the *ADH1B* locus.

**Fig 5.**
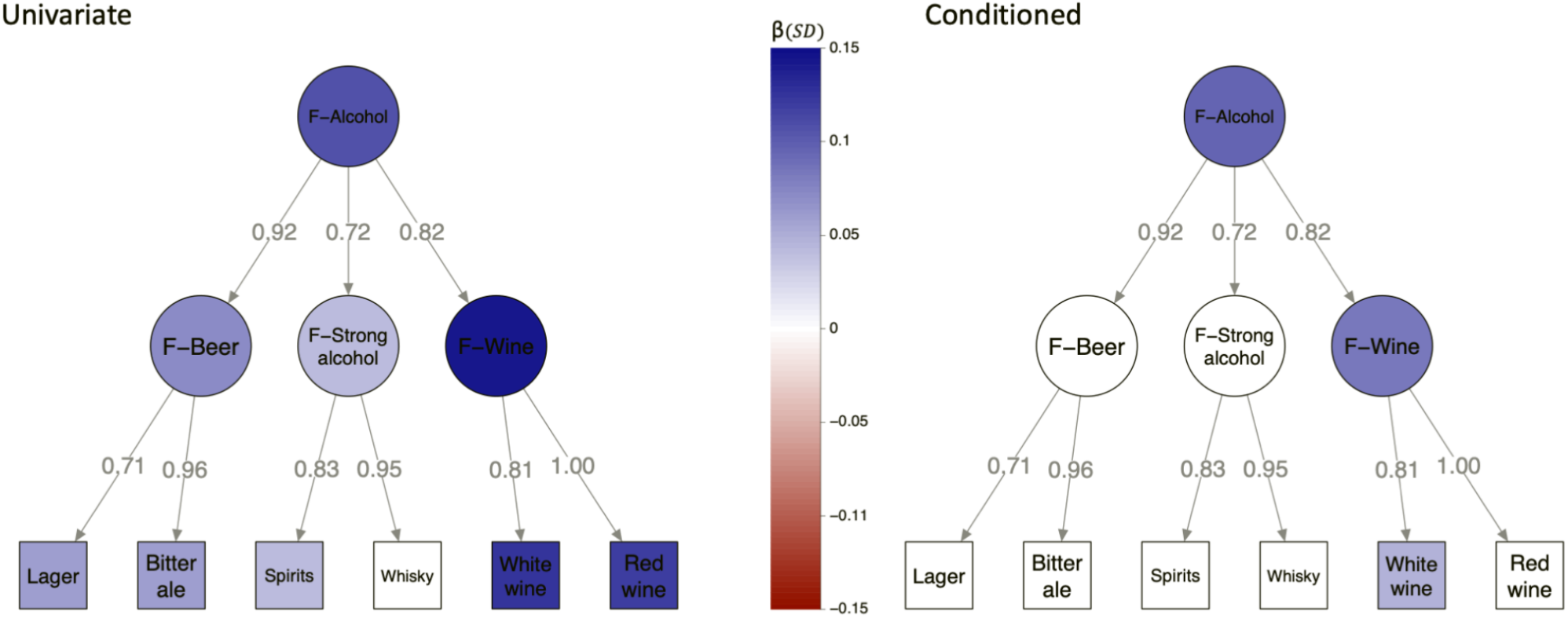
Example of univariable vs conditioned analysis of rs1229984. The path graph represents the hierarchical model up to the alcohol trait. Numbers over the edges report the standardised loadings. Colour is proportional to effect size. Effect sizes with p<1.4×10^-3^ have been shrunk to 0.

As can be seen, there was strong evidence that the rs1229984 SNP was associated with most alcoholic drinks. However, this SNP had a lesser effect on the stronger alcoholic drinks, suggesting a different weight of alcohol-liking, depending on its concentration. After the conditional analysis, only the effect of rs1229984 on alcohol remained unchanged, suggesting that *ADH1B* may exert most of its effect on alcoholic beverages through liking of alcohol in general, although residual effects remain on wine and white wine. Figures for most likely causal SNPs of the 208 association clusters comprising the full model can be found in Supplementary File 3 and Supplementary Table 10

### Tissue and Functional enrichment analysis

Functional enrichment expanding the gene selection to all loci with p<5×10^- 8^ (Supplementary Table 3), resulted in very strong enrichment of cellular components and biological processes related to neurons and specifically to glutamatergic and GABAergic synapses (Fig 3), both important and well-known modulators of hedonic responses to foods. These results are in line with the tissue enrichment analysis, where the only tissue that showed evidence for upregulation was the brain (Fig 6; Supplementary Table 11- 12)

**Fig 6.**
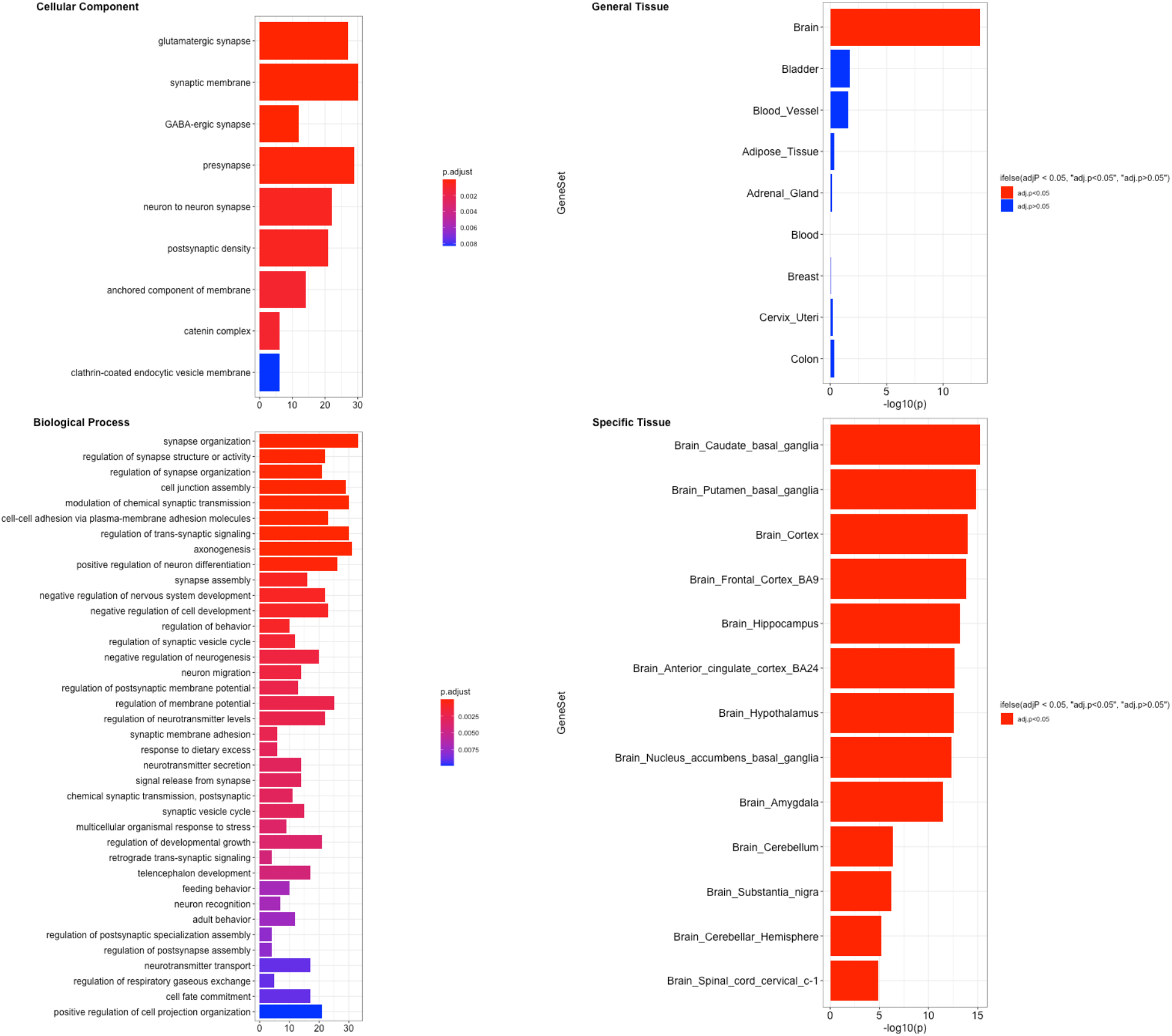
Enrichment analysis of food-liking genes. Figure represents the results of the GO terms and tissue up-regulated genes using the prioritised genes from all loci with p<5×10^-8^. Right panels show the summarised significant GO Terms (FDR <0.05) while the left ones report the tissue enrichment using the general tissues (upper panel) and the specific ones (bottom panel).

### Genetic correlation with brain morphology and connectivity traits

Genetic correlations with the brain morphology traits and IC100 rfMRI networks (Fig 7 and Supplementary table 13) evidenced clear differences in both types of traits. The morphological associations with the learned and low-caloric liking dimensions are characterized by negative correlations with cortical thickness in frontal (middle frontal, inferior frontal and orbital), parietal (intra-parietal and pre-cuneus) and occipital (cuneus, calcarine and lateral) areas, as well as positive correlations with cortical surface area in frontal/parietal transition area at the base of the (peri) central sulcus, in the temporal lobe in the fusiform area, and insula. In contrast, the Highly palatable liking dimension shows negative correlations with striatal volumes (in putamen and caudate) and no evident positive correlations.

**Fig. 7.**
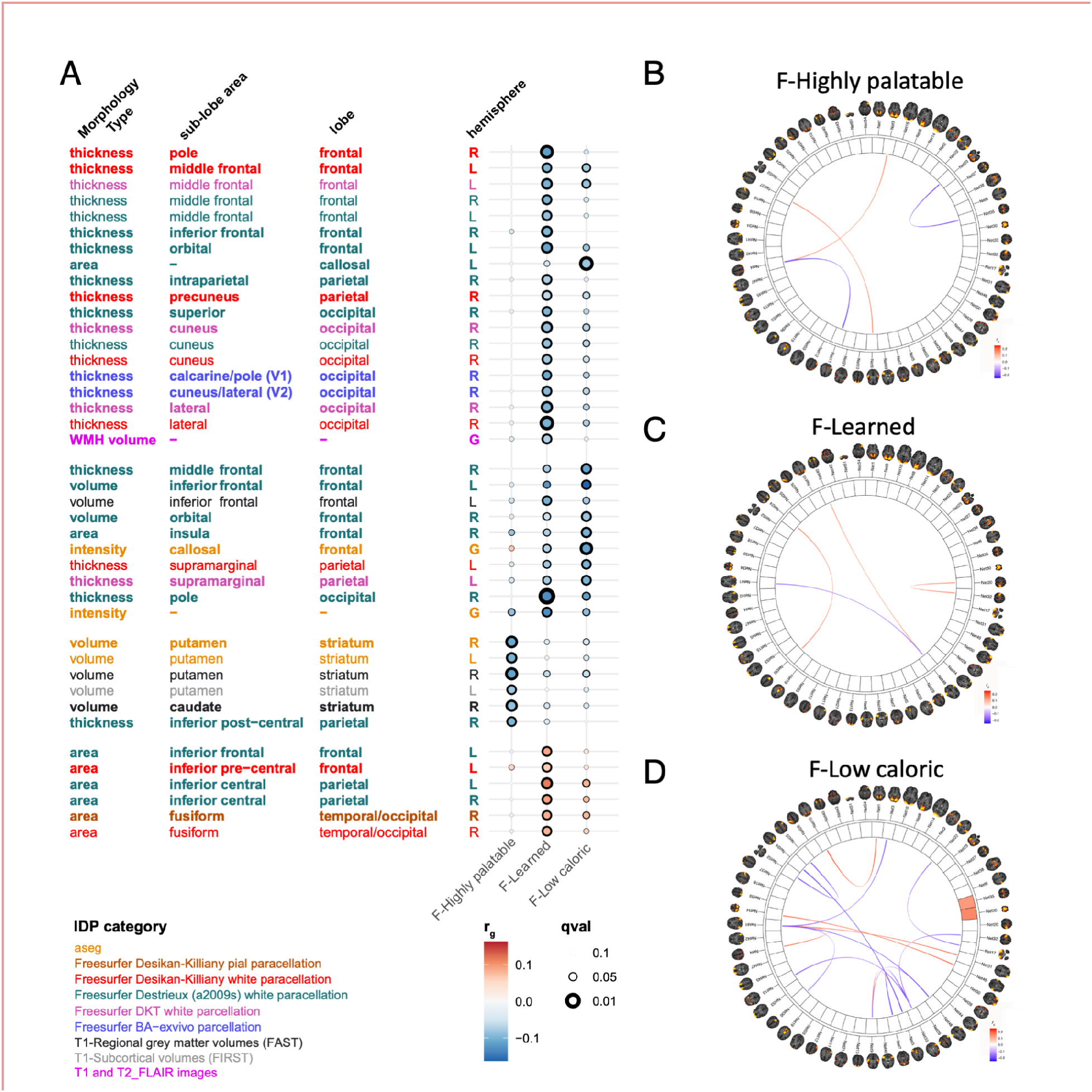
Genetic correlations between three main food-liking dimensions and brain MRI traits. Only traits with qvalue<0.05 have been reported. Panel A reports the genetic correlations between the three main liking dimensions and brain MRI morphological traits. Colour reflects the atlas used while size of the dots size is proportional to q-values. Panels B,C and D genetic correlations with the ICA100 network traits.

The connectivity network trait associations are also characterized by overlap in networks between learned and low-caloric, which both show (positive and negative) associations with frontal (somato-motor, language), parietal (intra-parietal), temporal (hippocampus, fusiform) and occipital (cuneus) areas. The Highly palatable food liking dimension shows few associations with connectivity networks, and when it does, they are characterized by positive associations with rostral frontal-parietal networks in frontal eye fields and intra-parietal cortex.

Summarizing, the morphological and network connectivity associations of the food-liking dimensions show parallel effects in the brain, such that both learned and low-caloric factors show associations with morphology in frontal, parietal and occipital areas and connectivity in networks involving the same areas, while the high-palatable dimension shows distinct associations, notably a negative association with morphology of striatal areas.

## Discussion

In this work, we have for the first time examined the genetic bases of food liking in a wide and comprehensive way. We have shown that it is possible to use genetic correlations to study the relationships between the food traits highlighting the complexity of these relationships and identifying three main distinct overall dimensions. We have also shown that these dimensions show different correlation patterns with both morphological and functional brain MRI traits. Furthermore, we have identified 171 loci involved in 1401 locus-trait associations, most of which have never been described before. Finally, we have used genomic structural equation modelling to disentangle many of the associations highlighting the main effects from those at least partly mediated through the effect of other food traits.

Food liking has been consistently shown to be a heritable trait in twin studies^11–16^. Here, we have shown that food liking also has a non-negligible SNP heritability and that it is twice as big as that of food consumption, in line with the idea that food liking is more influenced by biology than actual behaviour.

The fact that the genetic correlations between liking and food behaviour was relatively high, even when measured ∼10 years apart, suggests that the genetic factors underlying these two processes are very similar, while differences likely arise mostly from environmental factors and from the inherent differences between liking and choice. The fact that food liking is still so strongly correlated to consumption, even if measured later in time, suggests that food liking is relatively stable through time, at least in adults. Looking at the comparison between genetic and phenotypic correlations amongst the food items, they resemble each other quite closely (r=0.91), although the genetic correlations are twice as big as the phenotypic correlations. This likely reflects the random measurement error inherent in the use of questionnaires in measuring food liking and shows that genetic correlations may have advantages to assessing inter-relationships among food-related phenotypes. This strong relationship has been particularly useful in defining our hierarchical model, increasing our ability to identify the underlying dimensions common to multiple foods.

Whilst the current study is not the first to map how liking for different foods are related to each other, this is the largest and most comprehensive study to date, having used more than 150 thousand people and covered a wide range of food groups and flavours. In many cases, foods were clustered as expected (e.g., fresh vegetables and fruit) but in other cases have highlighted big differences in foods which are commonly considered as a single group. For example, while the genetic correlation between “cooked vegetables” and “salad vegetables” is very strong (0.79), when we consider also vegetables with stronger tastes such as spinach or asparagus (the “strong vegetables” group), this results in a much weaker correlation (0.38 and 0.54, respectively), despite the fact that these items would have generally all been considered “vegetables”. Our hypothesis-free approach thus captured these previously undescribed differences, which are of great importance in interpreting the results of nutritional studies.

When compared with the results from Vink et al^15^, our results show a clear resemblance between our first order traits and those identified through PCA. However, our strategy of using a multi-order hierarchical model allowed the identification of only a few higher order dimensions, highlighting the minimal correlation between very high reward foods such as sweets, meat and fried (the “F-Highly palatable” group) and other lower caloric and stronger taste intensities (F-Low caloric and F-Learned).

Looking at the genetic correlation with other complex traits we can see that the F-Highly palatable factor is, as expected, correlated with a worse anthropometric and lipid profiles, with signs of a diet rich in protein and salt. The F-Low caloric and F-learned show the opposite pattern, both associated with lower indices of obesity and a better blood lipid profile, with a diet lower in salt and protein. When we however look closer, these two factors do show some differences. The F-Learned factor is associated with a higher educational attainment and a sedentary job, likely indices of higher socioeconomic status, while for the F-Low caloric we see a different pattern where there is no correlation with educational attainment but a positive one for non-sedentary jobs.

Looking at the genetic correlations with the brain MRI morphological traits, while F-low caloric foods and F-Learned ones again show some agreement, the F-high palatable foods shows none with the other liking dimensions. Strikingly the Highly palatable foods correlated only (negatively) with striatum in putamen and caudate. Over-consumption of highly palatable energy-dense foods and adiposity are both associated with downregulation of neural responses in these areas. When we look at the areas involved with the other two other dimensions, we note they associate with areas involved with sensory responses, identification and decision making. These results indirectly confirm and validate our findings showing that the dimensions we have derived are not just an artefact of statistical inference, but correspond to true biological processes. Alternatively, they may reflect adaptations to dietary choices that result from the liking dimensions. They also suggest the existence of two distinct processes, mostly independent from each other, which underlie liking for the two groups of foods. This has profound implications in how food preferences in these two domains arise and in the shaping of future studies aimed at understanding them better.

Many studies which have looked at the genetics of food liking have focused on taste receptors, particularly on bitter ones. In this study, we have been able to confirm some of the previous findings such as that of the *TAS2R43- 46* locus and coffee liking^21^. For example, we observed a strong association between *TAS2R38,* responsible for PROP and PTC bitter taste, and both alcoholic beverage and salt liking, confirming our and others’ previous results on consumption^17^. We could not, however, replicate the association with any vegetable and, in fact, we found only weak evidence for such an association with broccoli, which was also in the opposite direction of what would be expected considering previous candidate gene studies. Given that we have looked at a large range of vegetables and the large sample size used, this result questions all previous candidate gene studies that have identified such associations^46^. Similarly, we found little evidence for an association with any of the genes coding for the sweet and umami receptor subunits (*TAS1R1-3*), again questioning some previous reports in much smaller samples of the association between these genes and sweet liking^46^.

When we look at the genes associated with flavour perception (see Fig S2), namely taste and olfactory receptors, we found that they associate only with the learned and low caloric foods and never with the Highly palatable foods. It is possible to speculate that this may have an evolutionary meaning, where variants which would lower liking of caloric dense foods such as those in the Highly palatable foods would be selected against, while those which increased acceptance of learned foods which are generally more aversive, would expand one’s diet and thus chances of survival. Further, more specific evolutionary genetics studies are needed to test this hypothesis.

Many genes already known to be associated with the consumption of specific foods showed a more complex association pattern, influencing a much broader range of food likings. For example, we have found that the variant rs1229984 within the *ADH1B* gene was expectedly associated with liking alcoholic beverages, mirroring the results on alcohol consumption. However, when we looked beyond simple genome-wide significance and reduced our p-value threshold, we found that it shows a marginal association with liking sweet foods with a concordant direction of association (see Fig S3). A recent GWAS of sweet liking^23^ conducted in a Japanese cohort where *ALDH2*, a variant known to be associated with alcohol consumption, is also associated with sweet liking but with the opposite effect where the allele associated with higher liking of alcohol is associated with lower liking of sweet foods. Both *ADH1B* and *ALDH2* gene products are responsible for metabolising alcohol in the liver and their association with alcohol consumption is believed to be through the accumulation of acetaldehyde, which gives an unpleasant feeling and thus will reduce alcohol consumption (and liking in our case) through conditioned learning. So although in both populations there is a genetic overlap between alcohol and sweet liking, this relationship is in opposite directions. These results suggest that the observed association is unlikely to be due to a biological mechanism but further studies involving people who have never consumed alcohol are needed to resolve this issue.

Another important example is *FGF21* which has been reported to be associated with consumption of sugar and protein^19,45^. Previous studies have shown that FGF21 is elevated by low protein and high carbohydrate consumption^47^. Soberg et al^48^ have previously shown that the rs838133 A allele is associated with lower levels of FGF21 and with higher consumption of sweet foods without an increase in energy intake or obesity. Our results are in line with these studies, with the A allele of rs838133 associated with higher liking of sweet foods, however when we look at the lower liked foods, although proteic foods are amongst them, they are represented by fish and cheese, but not by any of the meat traits (Fig S4). Moreover, we find a much wider range of traits which also include many strong-tasting vegetables and spices suggesting that the role of FGF21 is indeed to shift liking from sweet to savoury foods, but not necessarily all in the same way.

This example clearly shows how useful our results are in interpreting previous associations, greatly increasing our understanding of the phenomena behind food choices. Our results also highlight the importance of examining food liking as a whole instead of as sets of distinct sensations/food groups or macronutrients, where the interpretation of the results in one food dimension need to take account of the other factors in order to be properly interpreted. This is particularly important when studying the consequences of food liking on health status and particularly when performing Mendelian randomisation studies involving food traits.

Another interesting example is the association between a non-synonymous variant in the *GIPR* gene and liking of the foods in the low caloric group. *GIPR* encodes the receptor of glucose-dependent insulinotropic peptide (GIP), one of the two incretins and has been associated with BMI, in particular the A allele is associated with lower BMI^49^ and higher liking of low caloric foods and lower liking of fatty foods such as mayonnaise, cheese and cream (but not fatty meat products such as sausages) (Supplementary Fig. 5). GIPR encodes the receptor for the glucose-dependent insulinotropic peptide (GIP), that together with the Glucagon-like peptide-1 amide (GLP-1) represent the two human incretins. Amongst many other functions, incretins have been shown to regulate energy metabolism by acting in separate neuronal populations of the central nervous system^50^.

GLP-1and GIP have been shown to regulate food consumption synergistically by acting on the hypothalamic arcuate nucleus increasing neuronal activation and expression of pro-opiomelanocortin^50^. While both hormones are secreted in the presence of sugar, GIP responds also in the presence of free fatty acids^51^ . In a recent study^52^, CNS-*Gipr* knockout mice showed lower food intake when exposed to a high fat diet with smaller meals with consequent lower weight. Our results align very well, suggesting that GIPR, similarly to FGF21, is acting through a shift in preferences away from fatty foods and toward lower caloric foods, leading to a lower BMI. Both these examples point to regulation of food liking as a possible path through which to regulate food intake quality in order to, for example, help people comply with dietary plans beyond simple regulation of appetite.

In conclusion, we have presented the largest GWAS of food liking in more than 150 thousand individuals. We provided strong evidence that the dimensions of food liking are not only rooted in culture and familiarity but have an important biological basis, while identifying hundreds of novel associations between genetic variation across the human genome and liking of different foods. This not only greatly increases our knowledge in the field but opens up numerous paths for further studies aimed at better understanding the processes behind food choice.

## Supporting information

Diagram plots of the factor traits

Supplementary genetic correlation summary stats

## Acknowledgements

We would like to thank all the study participants without whom this study would not have been possible. We would like to thank, Prof Catherine Sudlow for her support to this project, Dr. Jo holyday for all the work during the final questionnaire definition and roll out. We would further like to thank Dr. Michel Nivard and Dr. Yakov A. Tsepilov for the advice and the great methods they have produced. This research has been conducted using the UK Biobank Resource under Application Number 19655.

The Viking Health Study – Shetland (VIKING): DNA extractions and genotyping were performed at the Edinburgh Clinical Research Facility, University of Edinburgh. We would like to acknowledge the invaluable contributions of the research nurses in Shetland, the administrative team in Edinburgh and the people of Shetland.

We are extremely grateful to all the families who took part in this study, the midwives for their help in recruiting them, and the whole ALSPAC team, which includes interviewers, computer and laboratory technicians, clerical workers, research scientists, volunteers, managers, receptionists and nurses. Genome-wide association data was generated by Sample Logistics and Genotyping Facilities at Wellcome Sanger Institute and LabCorp (Laboratory Corporation of America) using support from 23andMe.

We are very grateful to the municipal administrators of all INGI cohorts, for their collaboration on the project and for logistic support. We would like to thank all participants to this study.

## Author contributions

NP, EdG, KW, NJT, JFW, MGV, designed the study; CM, EdG,MM, DB,JFW, KW, PG provided/collected data; NP, NM, SMW,MM, EJG, KW, JJH, MGV,MPC analysed the data; EDG, DB JFW, PG, provided funding; NP, KW, NM, MGV, JFW, NJT,EDG, wrote the manuscript. All authors reviewed and provided comments to the text.

## Data availability

All GWAS results will be available through GWAS catalogue at the time of publication.

Supplementary file 3 can be downloaded at: https://drive.google.com/file/d/1wD92SjAQ0jGTQYaBRj8Dl9wxz_QZAIJo/view?usp=sharing

