## Supplementary Figs. 1-5 for "Large-scale genome-wide association study of food liking reveals genetic determinants and genetic correlations with distinct neurophysiological traits"

**Supplementary Figures**

Fig. S1 Comparison of genetic and phenotypic correlation between the food liking traits. Panel A shows the correlation plot for all the pairwise correlation between the food traits used in the analysis. Two main groups of foods are visible and have been highlighted with the black boxes. Panel B Scatterplot of all pairwise genetic and phenotypic correlations. The red line represents the regression line. Although the genetic and phenotypic correlations resemble each other very closely (r>0.9) genetic correlations are ~2x as big reflecting the large amount of noise present in the questionnaire data.

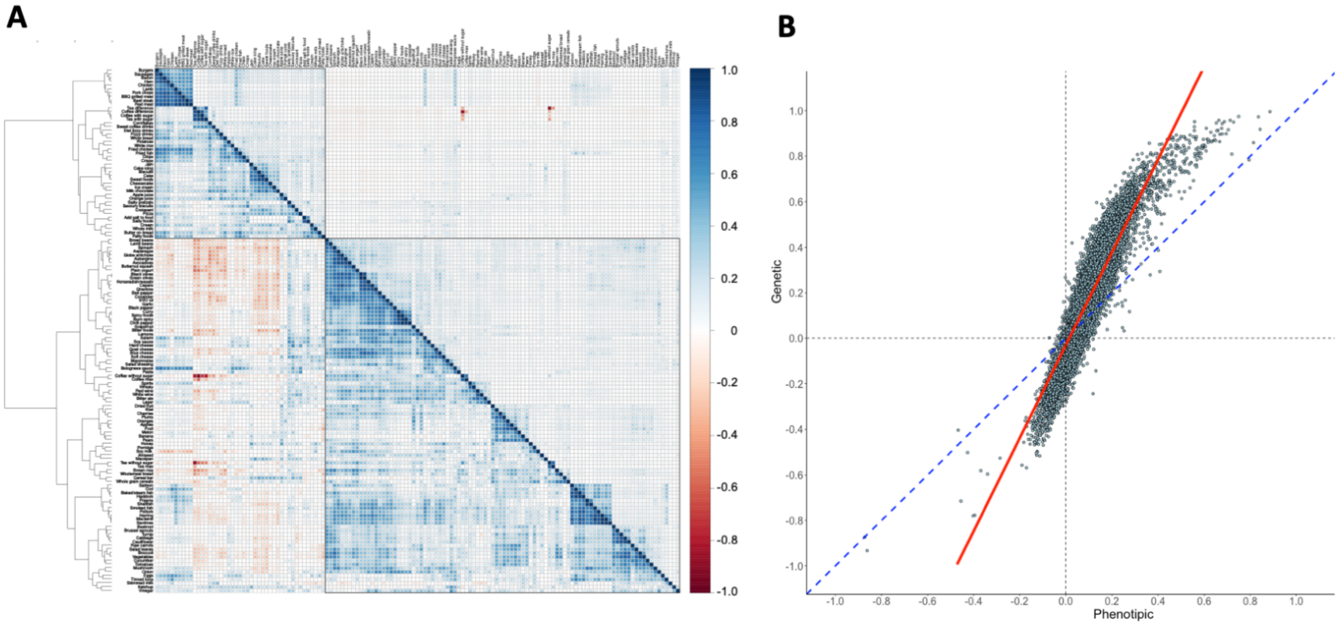

Fig. S2. Traits associated with flavour perception.
The figure highlights the traits associated to genes which encode taste or olfactory receptors. TRPV1 has been included as well as it is responsible for the burning sensation of capsaicin.

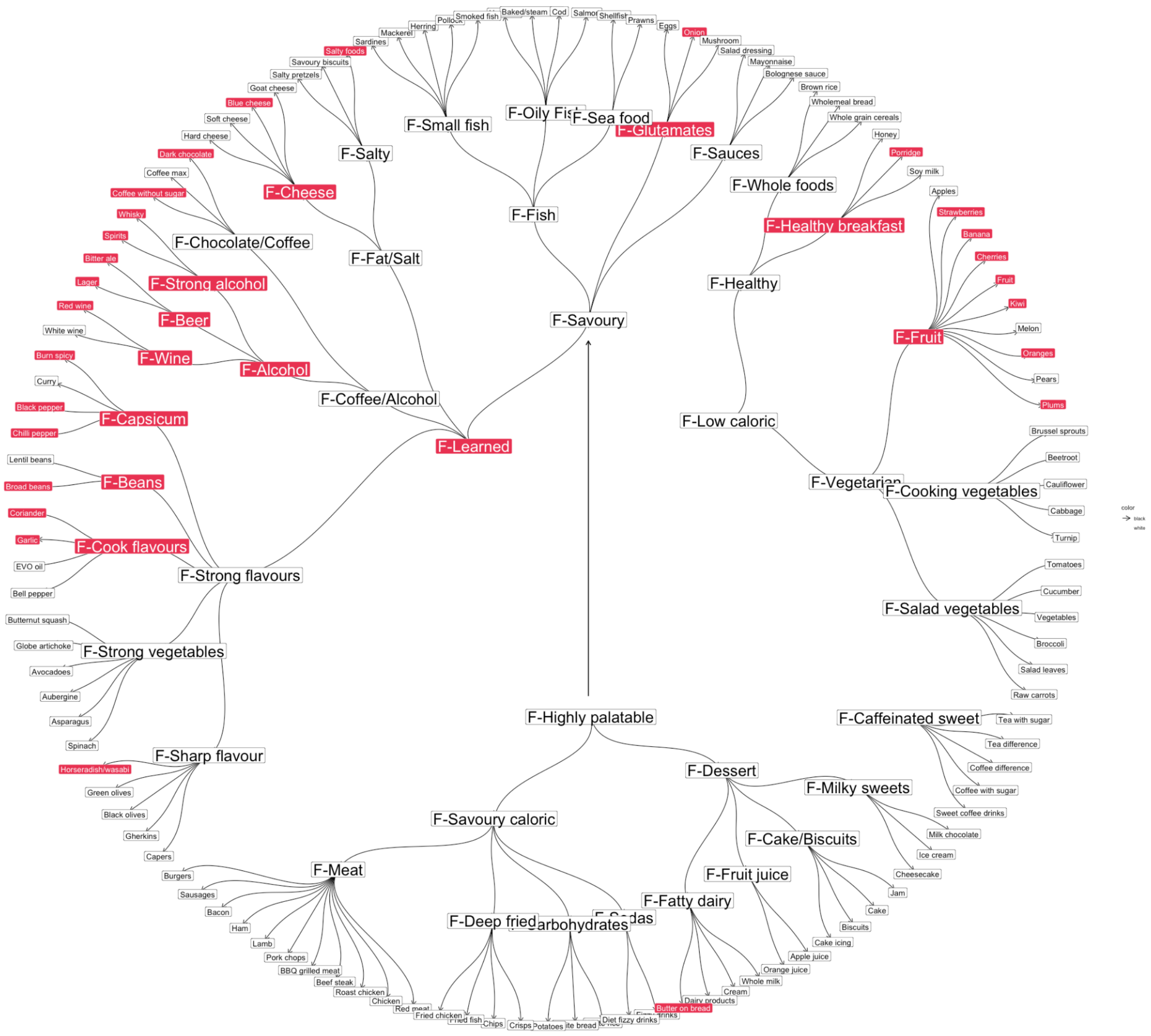

Fig S3. Effect of rs1229984 in *ADH1B* on food-liking traits. Forest plot of the effect of the C allele of rs1229984 on food liking. The point represents the point estimate while the bar represents the 95% CI. Only traits which colocalise and for which rs1229984 is the most likely causal SNP have been reported.

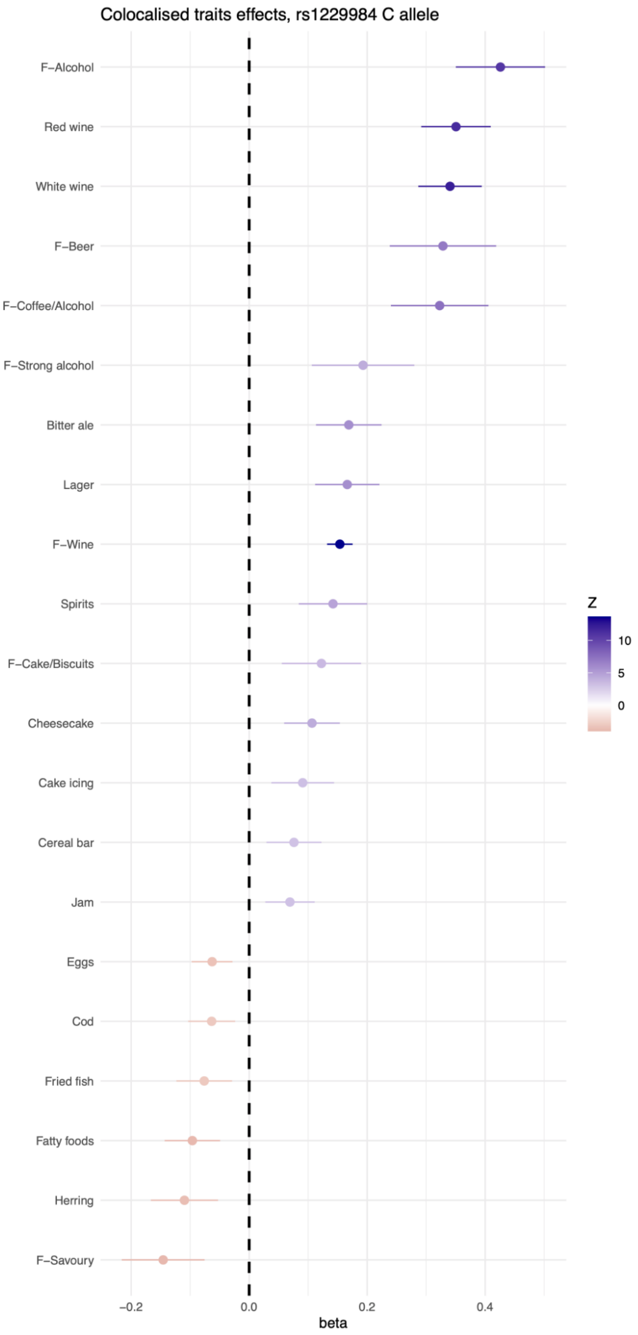

Fig S4. Effect of rs838133 in *FGF21* on food-liking traits. Forest plot of the effect of the A allele of rs838133 on food liking. The point represents the point estimate while the bar represents the 95% CI. Only traits which colocalise and for which rs838133 is the most likely causal SNP have been reported.

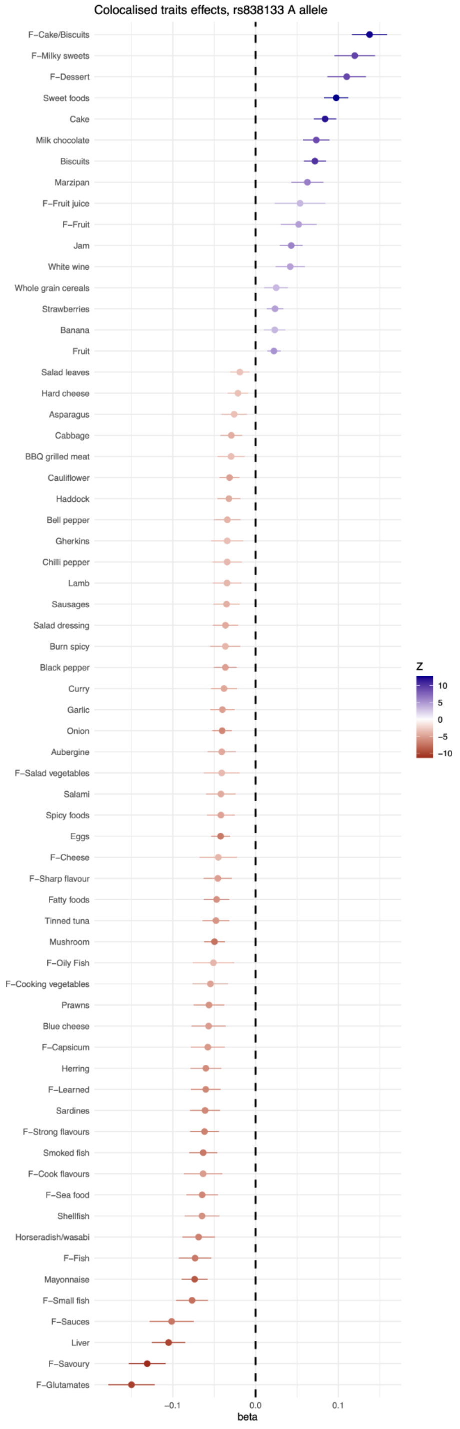

Fig S5. Effect of rs10423928 in *GIPR* on food-liking traits. Forest plot of the effect of the A allele of rs10423928 on food liking. The point represents the point estimate while the bar represents the 95% CI. Only traits which colocalise and for which rs10423928 is the most likely causal SNP have been reported.

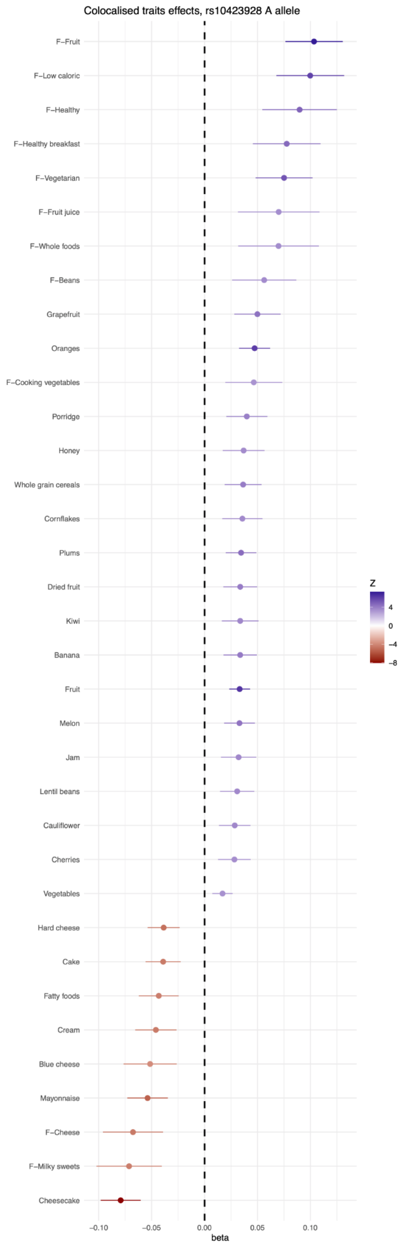
