## Supplementary material for "Large-scale genome-wide association study of food liking reveals genetic determinants and genetic correlations with distinct neurophysiological traits": Diagram plots of the factor traits

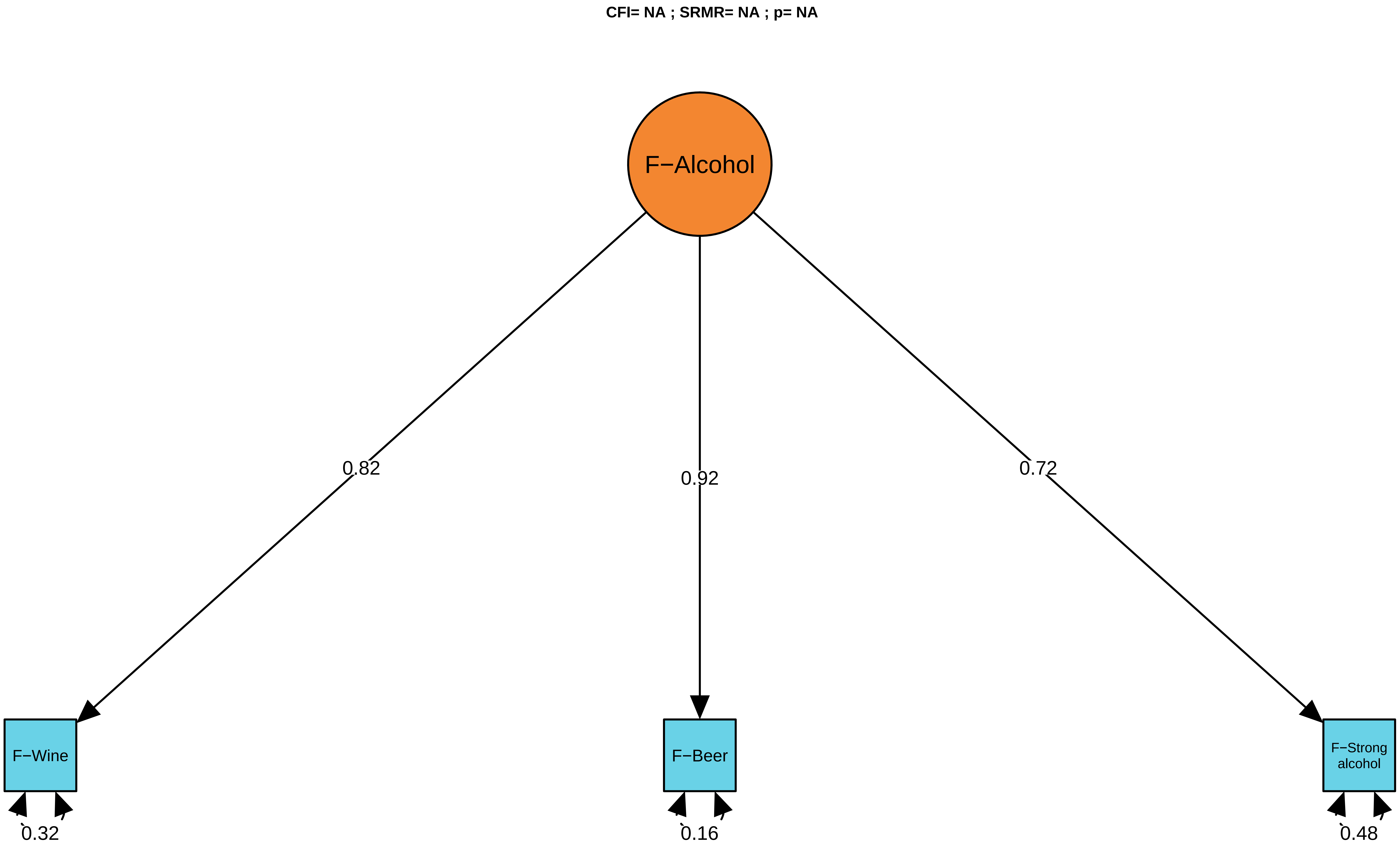

CFI= NA ; SRMR= NA ; p= NA
F−Alcohol
0.82
0.92
0.72
F−Wine
F−Beer
F−Strong alcohol
0.32
0.16
0.48

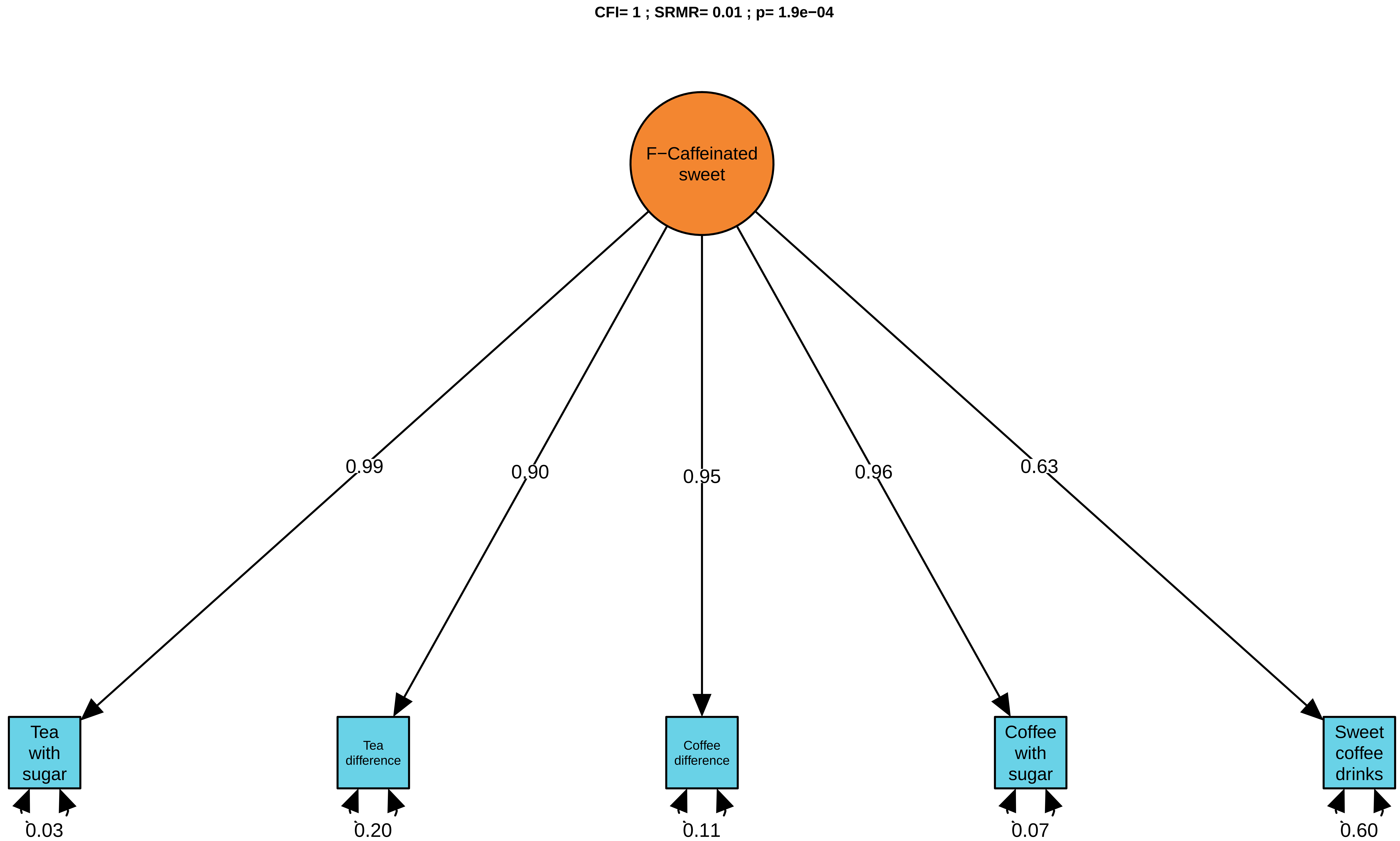
CFI= 1 ; SRMR= 0.01 ; p= 1.9e−04
F−Caffeinated sweet
0.99
0.90
0.95
0.96
0.63
Tea with sugar
Tea difference
Coffee difference
Coffee with sugar
Sweet coffee drinks
0.03
0.20
0.11
0.07
0.60

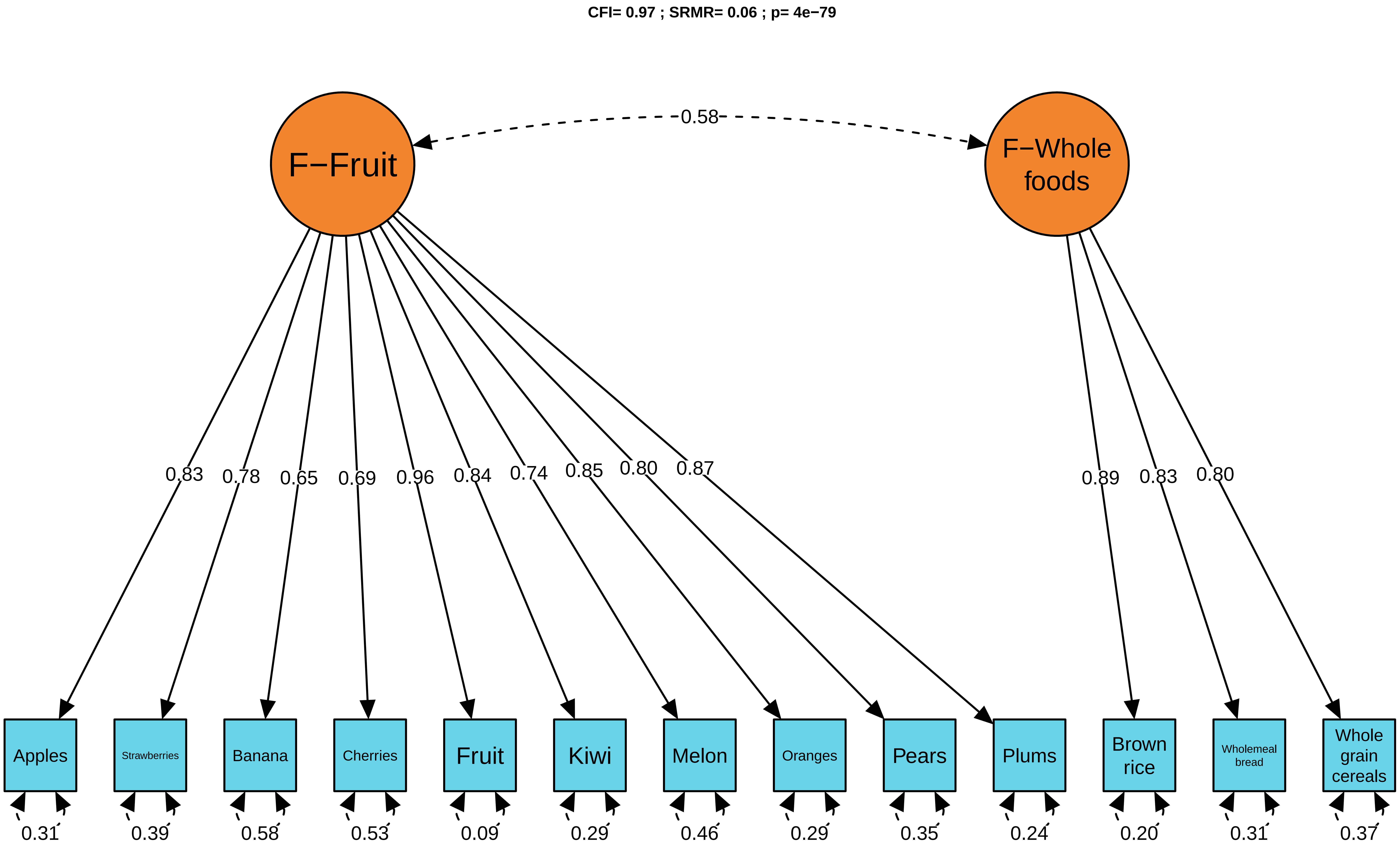
CFI= 0.97 ; SRMR= 0.06 ; p= 4e−79
F−Fruit
F−Whole foods
0.58
0.83
0.78
0.65
0.69
0.96
0.84
0.74
0.85
0.80
0.87
0.89
0.83
0.80
Apples
Strawberries
Banana
Cherries
Fruit
Kiwi
Melon
Oranges
Pears
Plums
Brown rice
Wholemeal bread
Whole grain cereals
0.31
0.39
0.58
0.53
0.09
0.29
0.46
0.29
0.35
0.24
0.20
0.31
0.37

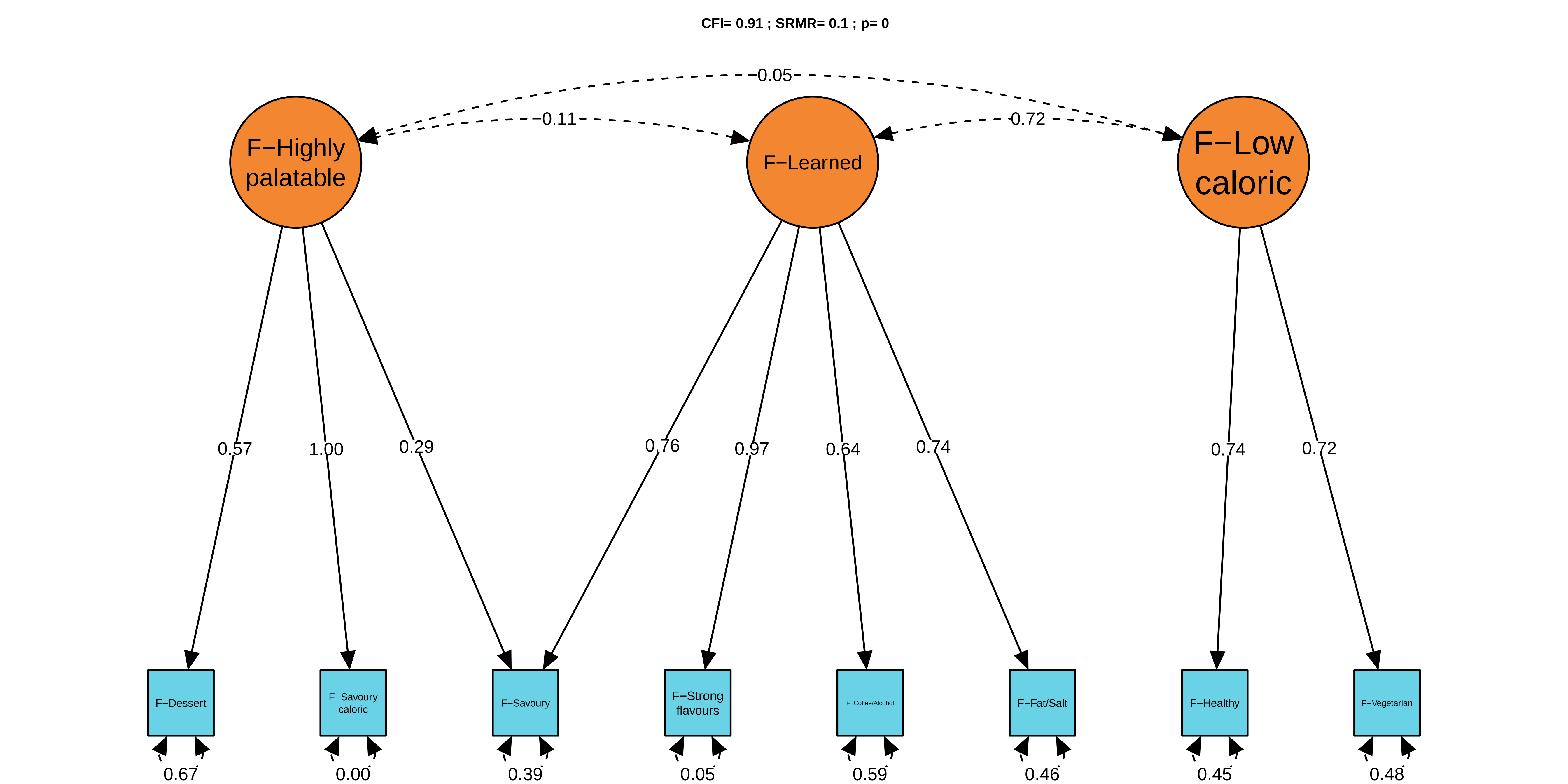

CFI= 0.91 ; SRMR= 0.1 ; p= 0
0.05
0.11
0.72
F−Highly palatable
F−Learned
F−Low caloric
0.57
1.00
0.29
0.76
0.97
0.64
0.74
0.74
0.72
F−Dessert
F−Savoury caloric
F−Savoury
F−Strong flavours
F−Coffee/Alcohol
F−Fat/Salt
F−Healthy
F−Vegetarian
0.67
0.00
0.39
0.05
0.59
0.46
0.45
0.48

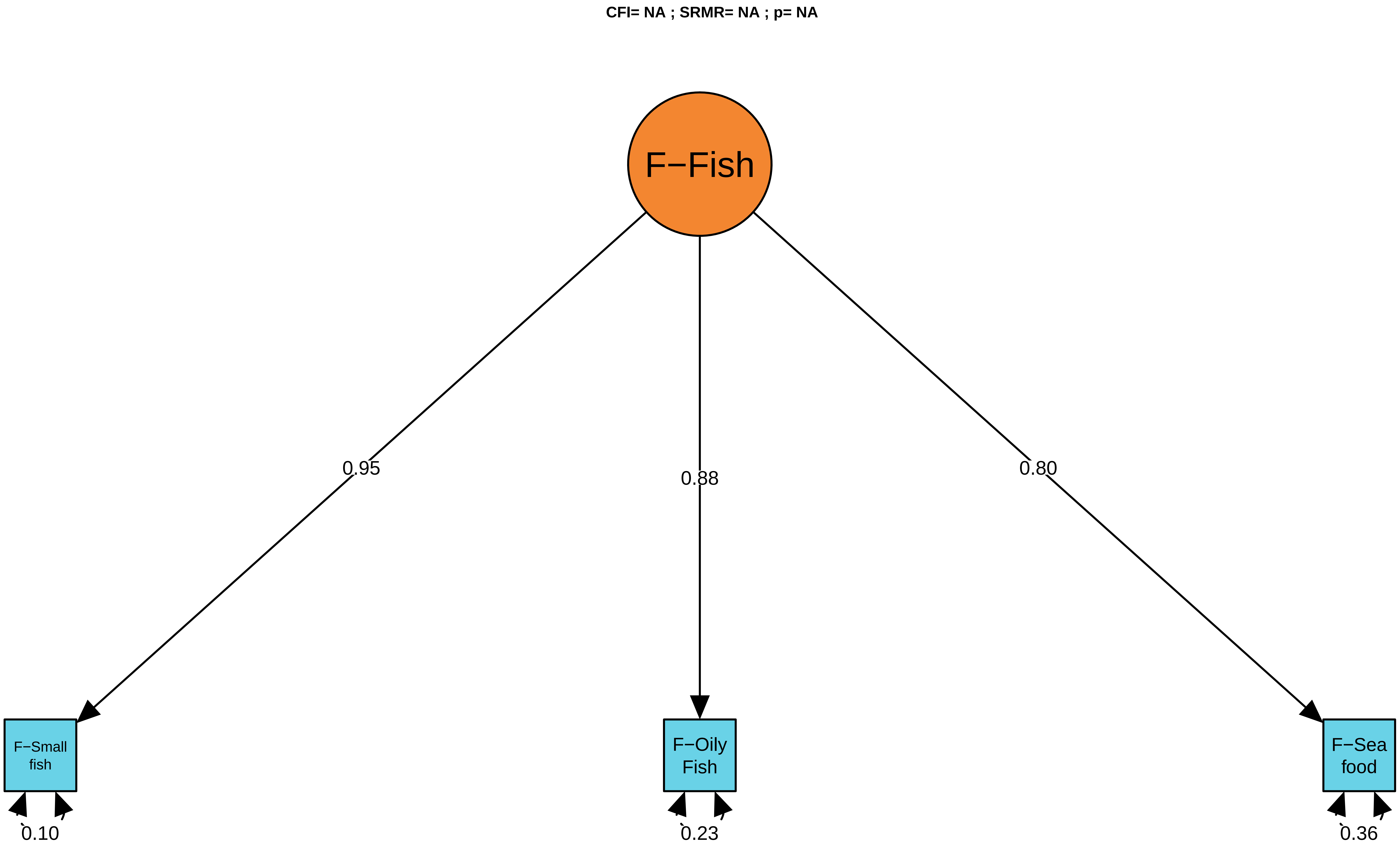
CFI= NA ; SRMR= NA ; p= NA
F−Fish
0.95
0.88
0.80
F−Small fish
F−Oily Fish
F−Sea food
0.10
0.23
0.36

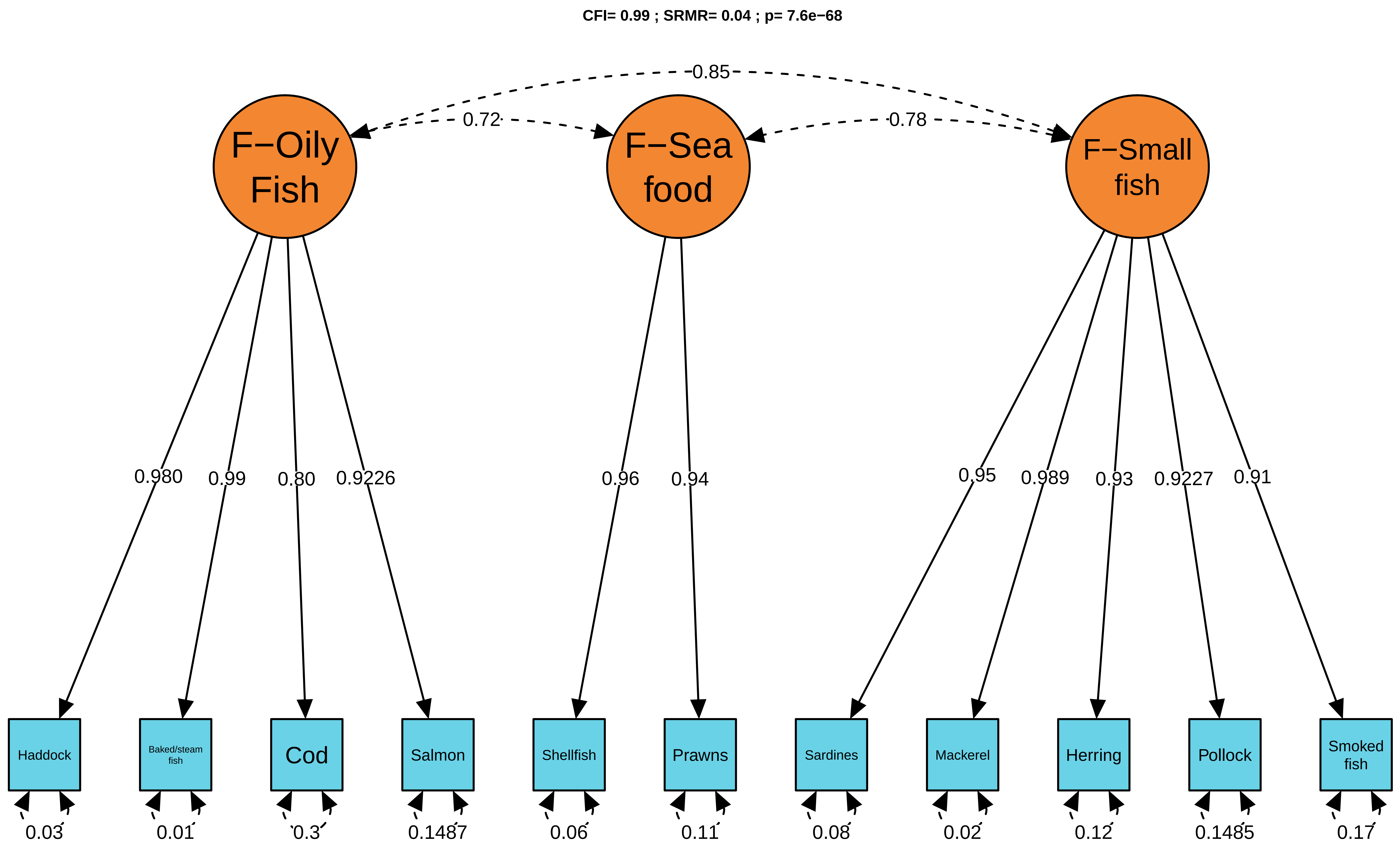

CFI= 0.99 ; SRMR= 0.04 ; p= 7.6e−68
0.85
0.72
0.78
F−Oily Fish
F−Sea food
F−Small fish
0.980
0.99
0.80
0.9226
0.96
0.94
0.95
0.989
0.93
0.9227
0.91
Haddock
Baked/steam fish
Cod
Salmon
Shellfish
Prawns
Sardines
Mackerel
Herring
Pollock
Smoked fish
0.03
0.01
0.3
0.1487
0.06
0.11
0.08
0.02
0.12
0.1485
0.17

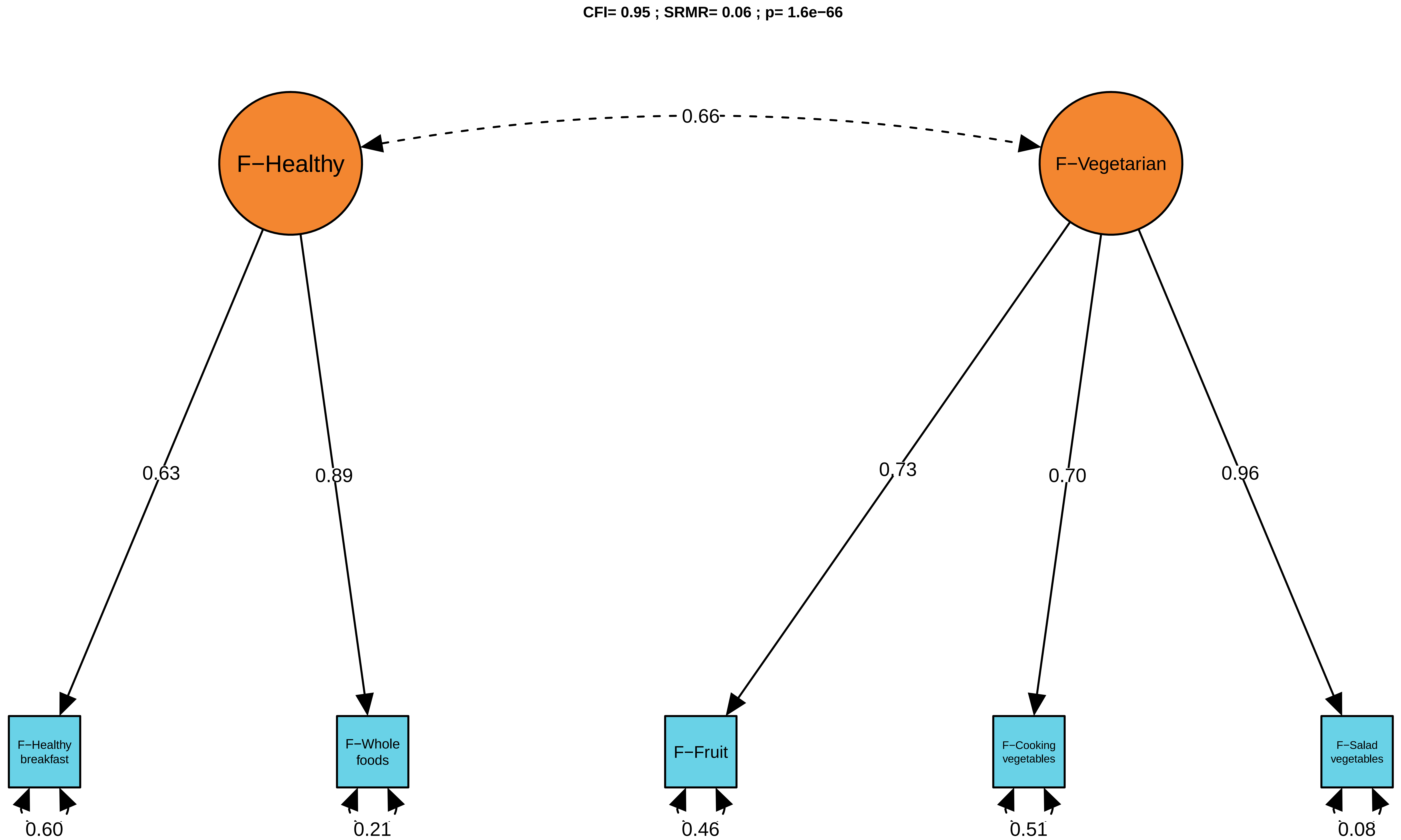
CFI= 0.95 ; SRMR= 0.06 ; p= 1.6e−66
F−Healthy
F−Vegetarian
0.66
0.63
0.89
0.73
0.70
0.96
F−Healthy breakfast
F−Whole foods
F−Fruit
F−Cooking vegetables
F−Salad vegetables
0.60
0.21
0.46
0.51
0.08

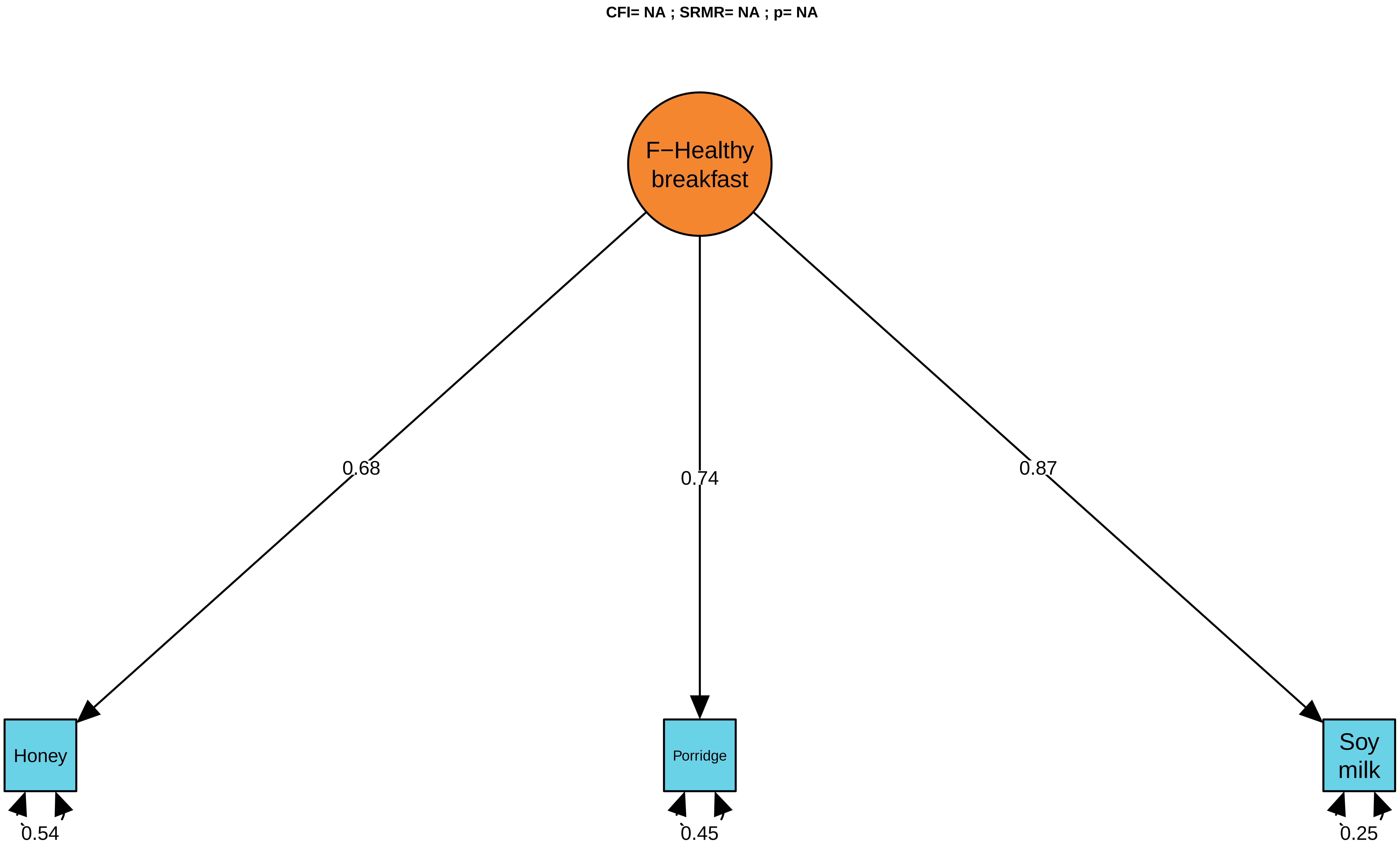
CFI= NA ; SRMR= NA ; p= NA
F−Healthy breakfast
0.68
0.74
0.87
Honey
Porridge
Soy milk
0.54
0.45
0.25

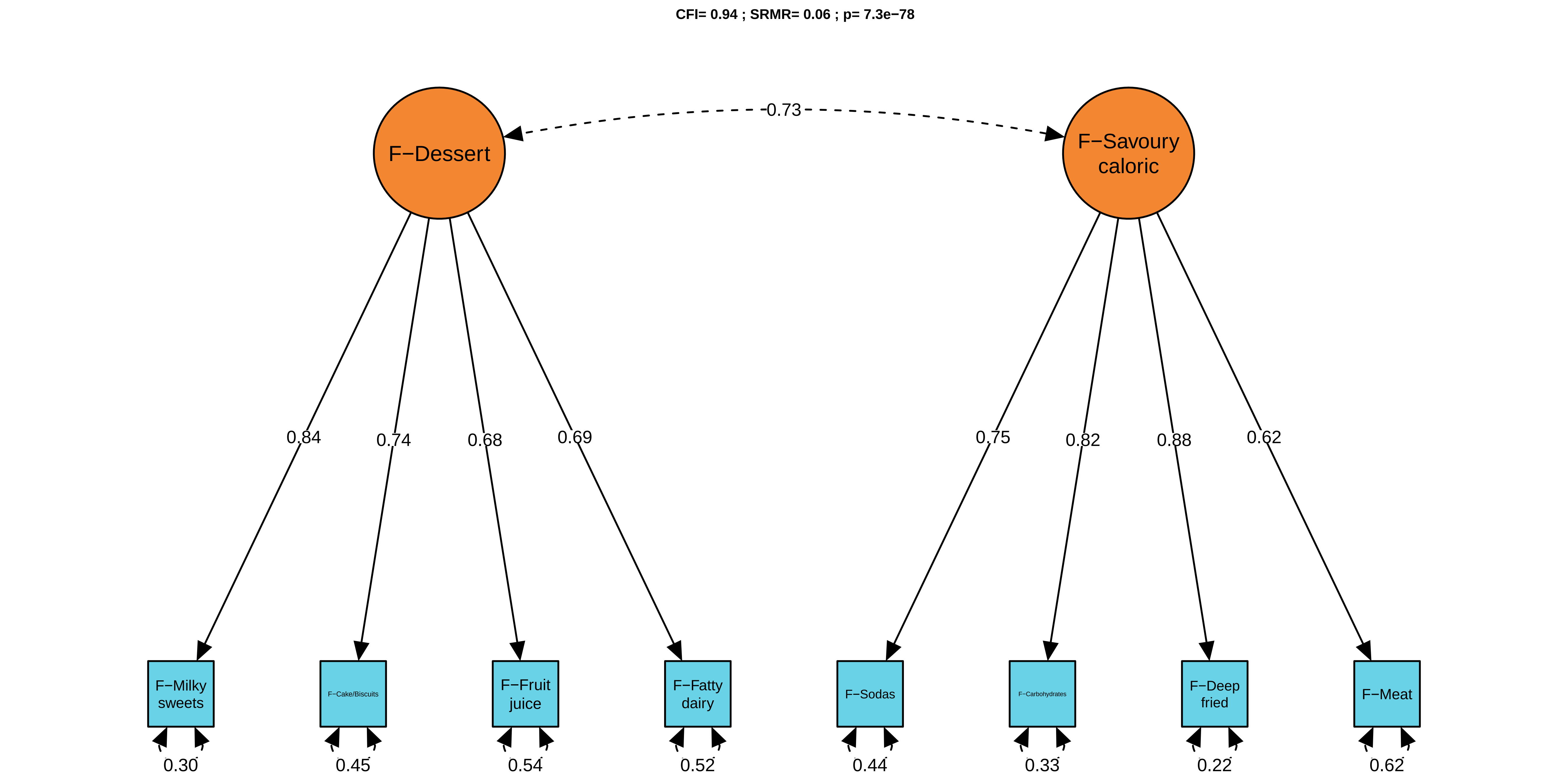
CFI= 0.94 ; SRMR= 0.06 ; p= 7.3e−78
F−Dessert
F−Savoury caloric
0.73
0.84
0.74
0.68
0.69
0.75
0.82
0.88
0.62
F−Milky sweets
F−Cake/Biscuits
F−Fruit juice
F−Fatty dairy
F−Sodas
F−Carbohydrates
F−Deep fried
F−Meat
0.30
0.45
0.54
0.52
0.44
0.33
0.22
0.62

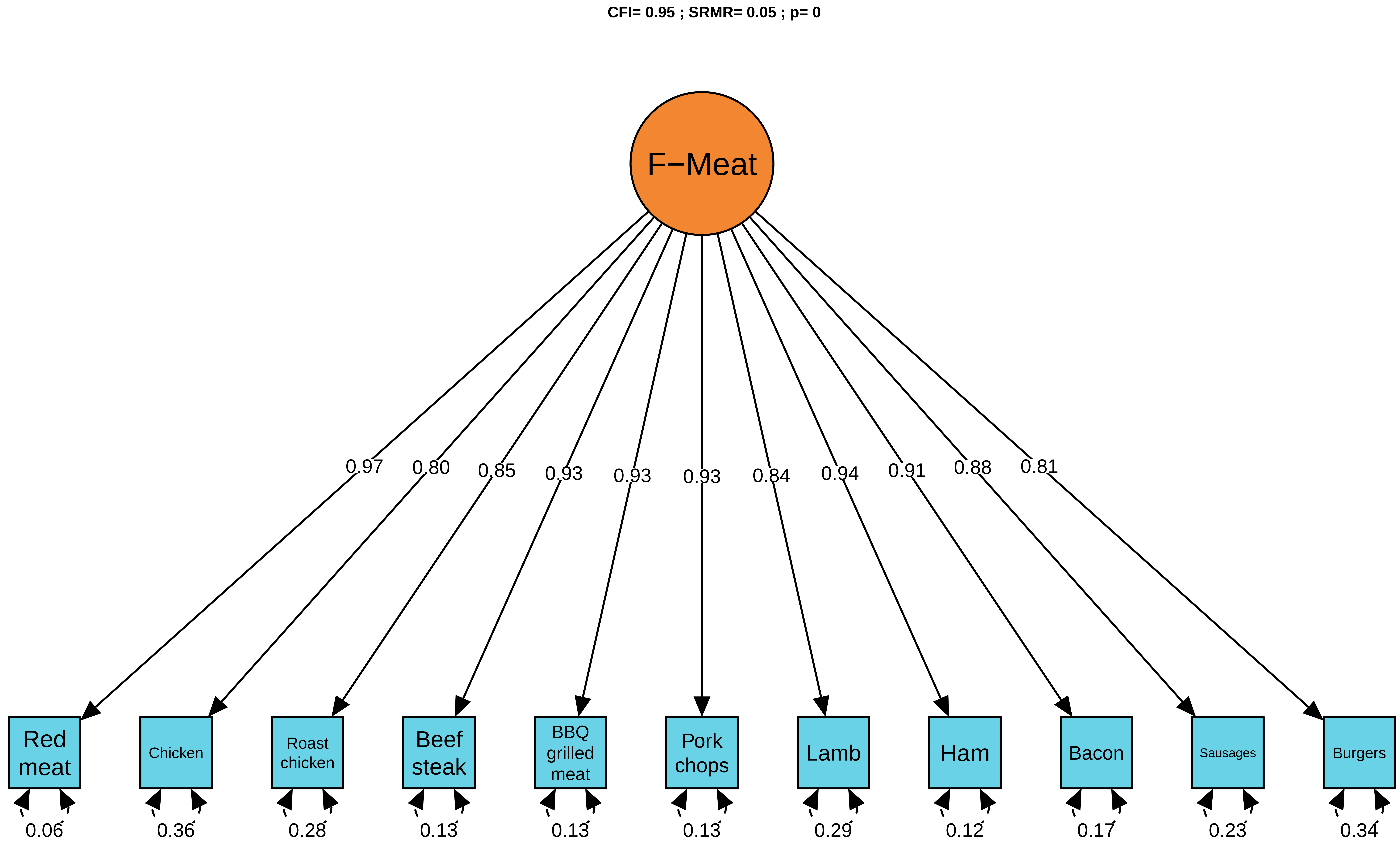
CFI= 0.95 ; SRMR= 0.05 ; p= 0
F−Meat
0.97
0.80
0.85
0.93
0.93
0.93
0.84
0.94
0.91
0.88
0.81
Red meat
Chicken
Roast chicken
Beef steak
BBQ grilled meat
Pork chops
Lamb
Ham
Bacon
Sausages
Burgers
0.06
0.36
0.28
0.13
0.13
0.13
0.29
0.12
0.17
0.23
0.34

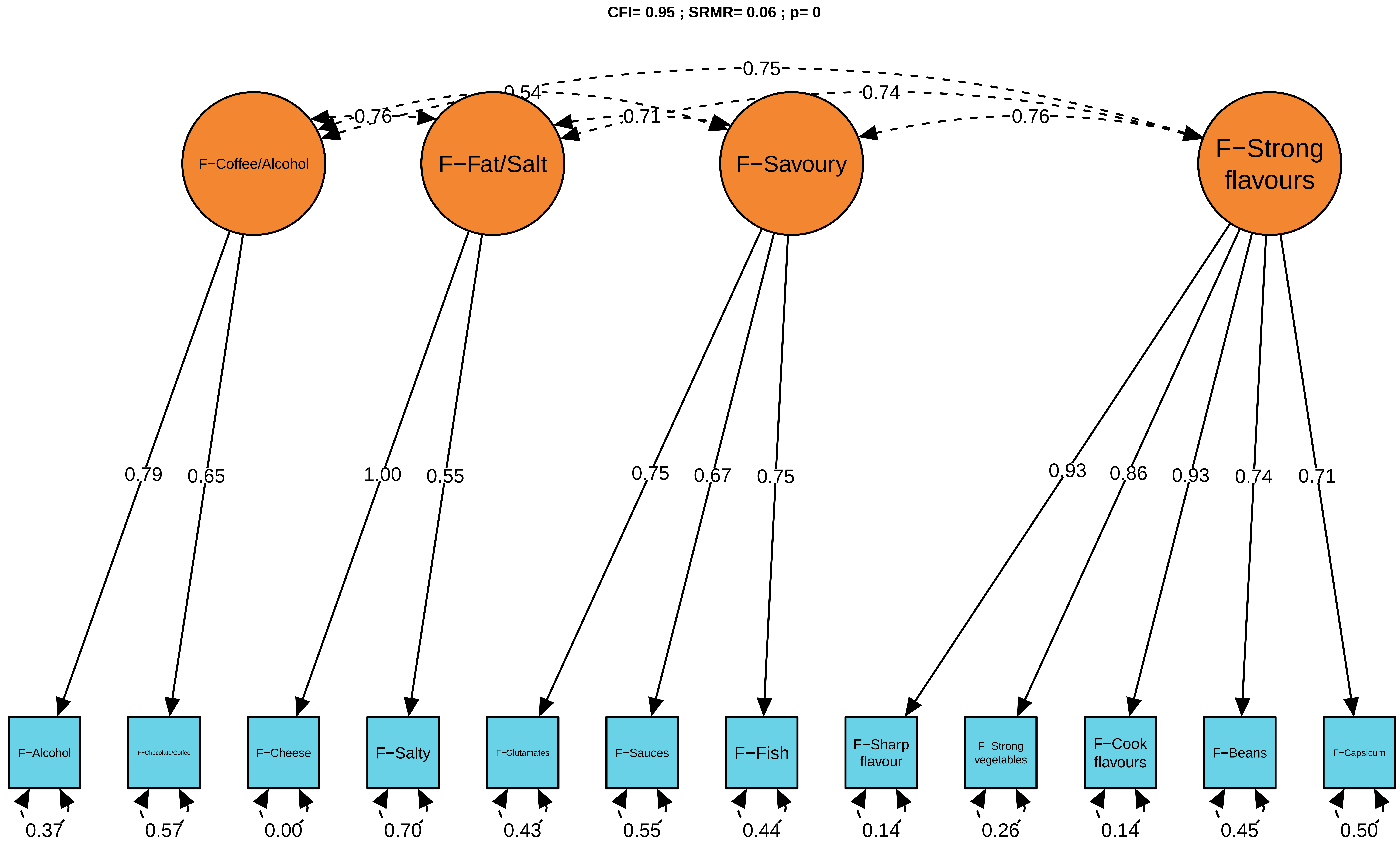

CFI= 0.95 ; SRMR= 0.06 ; p= 0
0.75
0.54
0.74
0.76
0.71
0.76
F−Coffee/Alcohol
F−Fat/Salt
F−Savoury
F−Strong flavours
0.79
0.65
1.00
0.55
0.75
0.67
0.75
0.93
0.86
0.93
0.74
0.71
F−Alcohol
F−Chocolate/Coffee
F−Cheese
F−Salty
F−Glutamates
F−Sauces
F−Fish
F−Sharp flavour
F−Strong vegetables
F−Cook flavours
F−Beans
F−Capsicum
0.37
0.57
0.00
0.70
0.43
0.55
0.44
0.14
0.26
0.14
0.45
0.50

**CFI= 0.98 ; SRMR= 0.05 ; p= 7.8e−22**

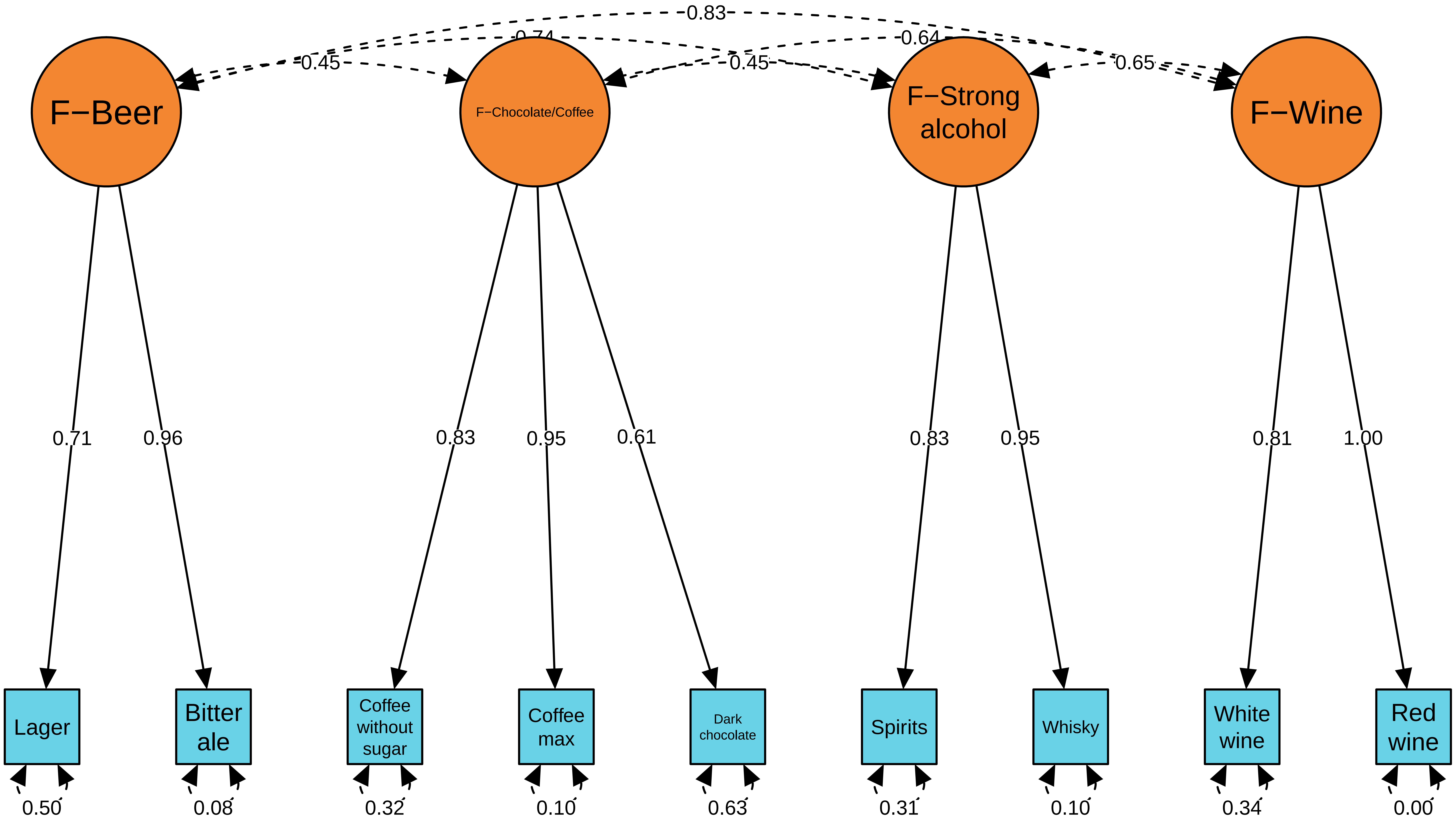

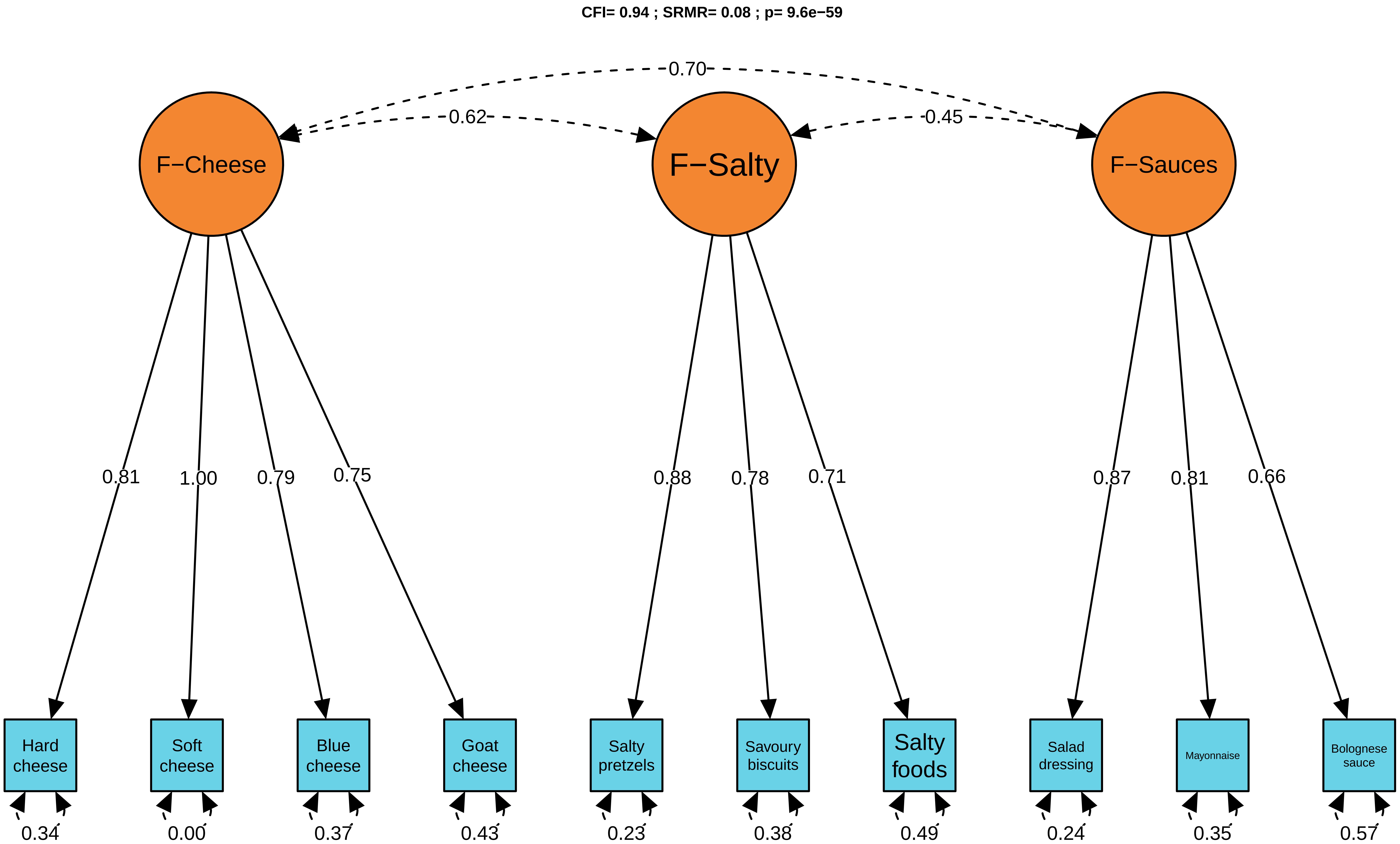

CFI= 0.94 ; SRMR= 0.08 ; p= 9.6e−59
0.70
0.62
0.45
F−Cheese
F−Salty
F−Sauces
0.81
1.00
0.79
0.75
0.88
0.78
0.71
0.87
0.81
0.66
Hard cheese
Soft cheese
Blue cheese
Goat cheese
Salty pretzels
Savoury biscuits
Salty foods
Salad dressing
Mayonnaise
Bolognese sauce
0.34
0.00
0.37
0.43
0.23
0.38
0.49
0.24
0.35
0.57

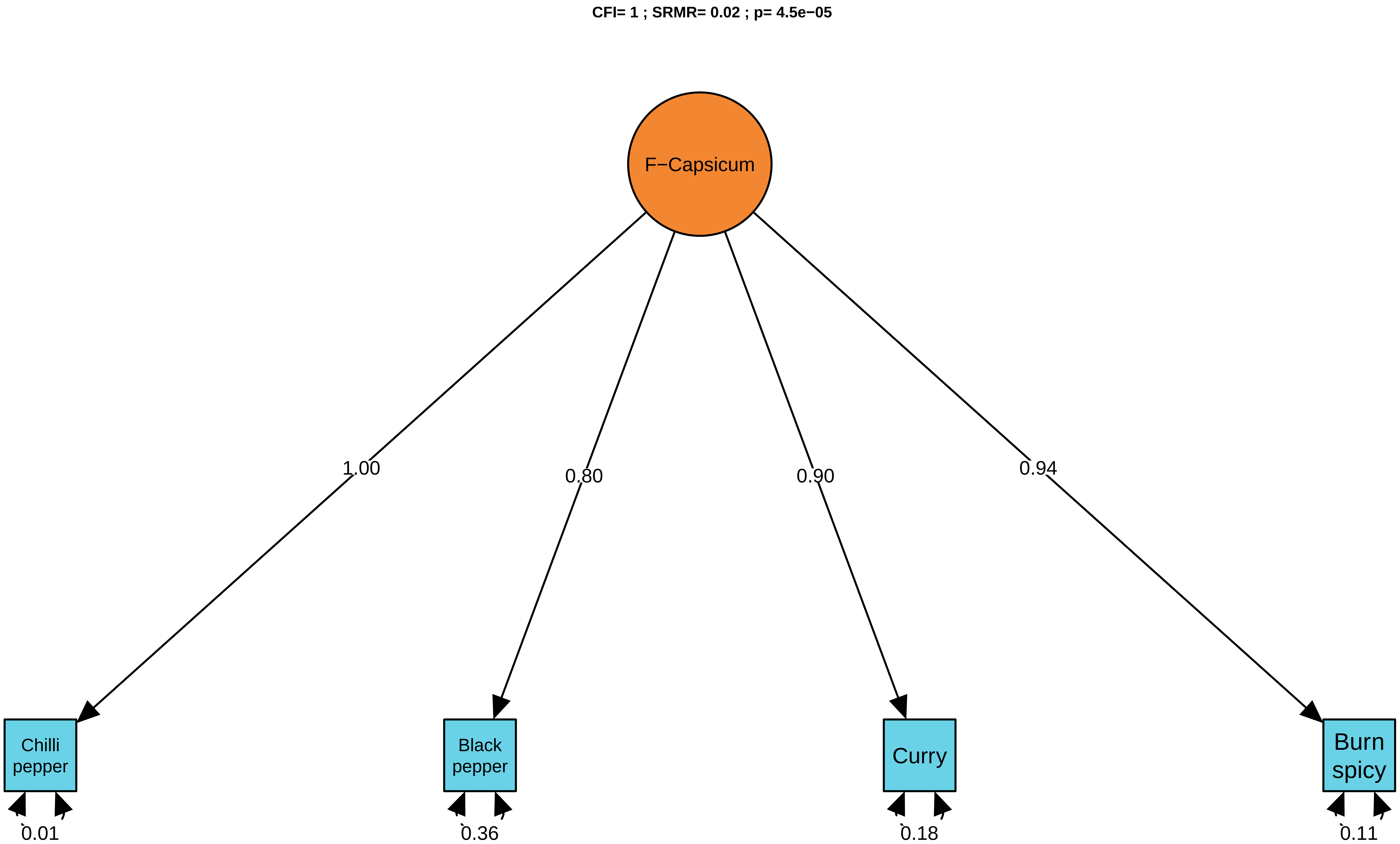
CFI= 1 ; SRMR= 0.02 ; p= 4.5e−05
F−Capsicum
1.00
0.80
0.90
0.94
Chilli pepper
Black pepper
Curry
Burn spicy
0.01
0.36
0.18
0.11

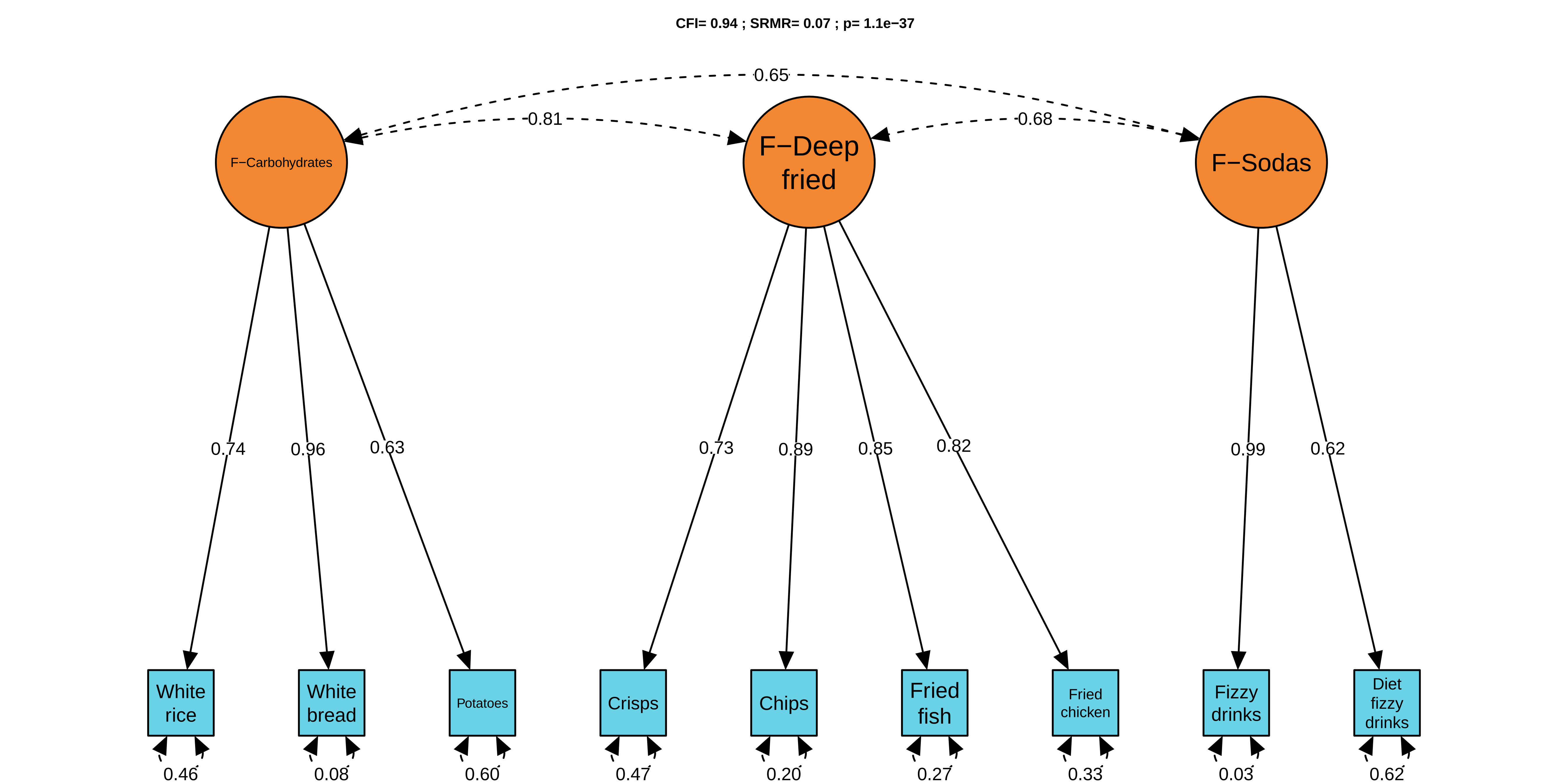

CFI= 0.94 ; SRMR= 0.07 ; p= 1.1e−37
0.65
0.81
0.68
F−Carbohydrates
F−Deep fried
F−Sodas
0.74
0.96
0.63
0.73
0.89
0.85
0.82
0.99
0.62
White rice
White bread
Potatoes
Crisps
Chips
Fried fish
Fried chicken
Fizzy drinks
Diet fizzy drinks
0.46
0.08
0.60
0.47
0.20
0.27
0.33
0.03
0.62

**CFI= 0.99 ; SRMR= 0.05 ; p= 0**

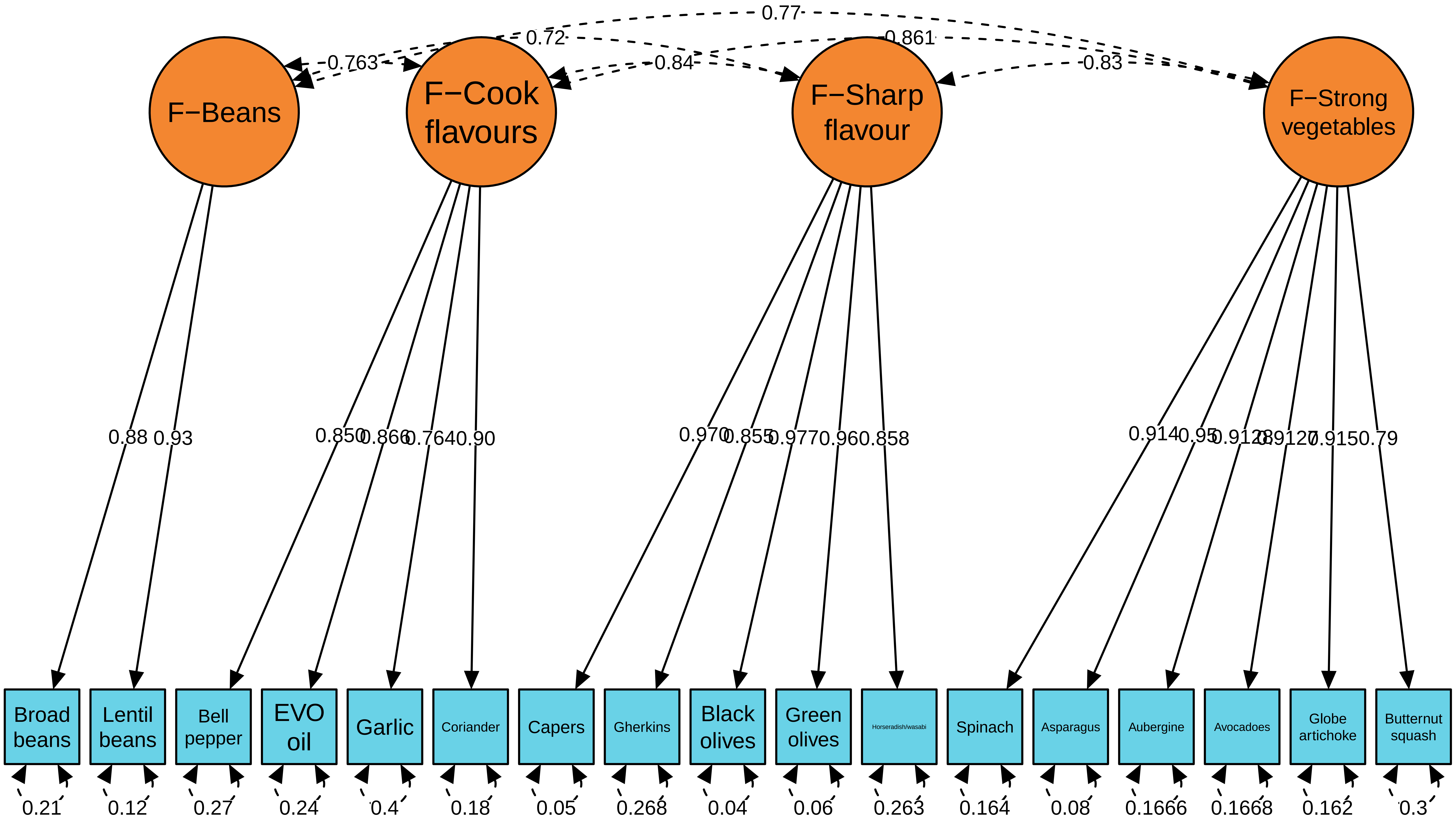

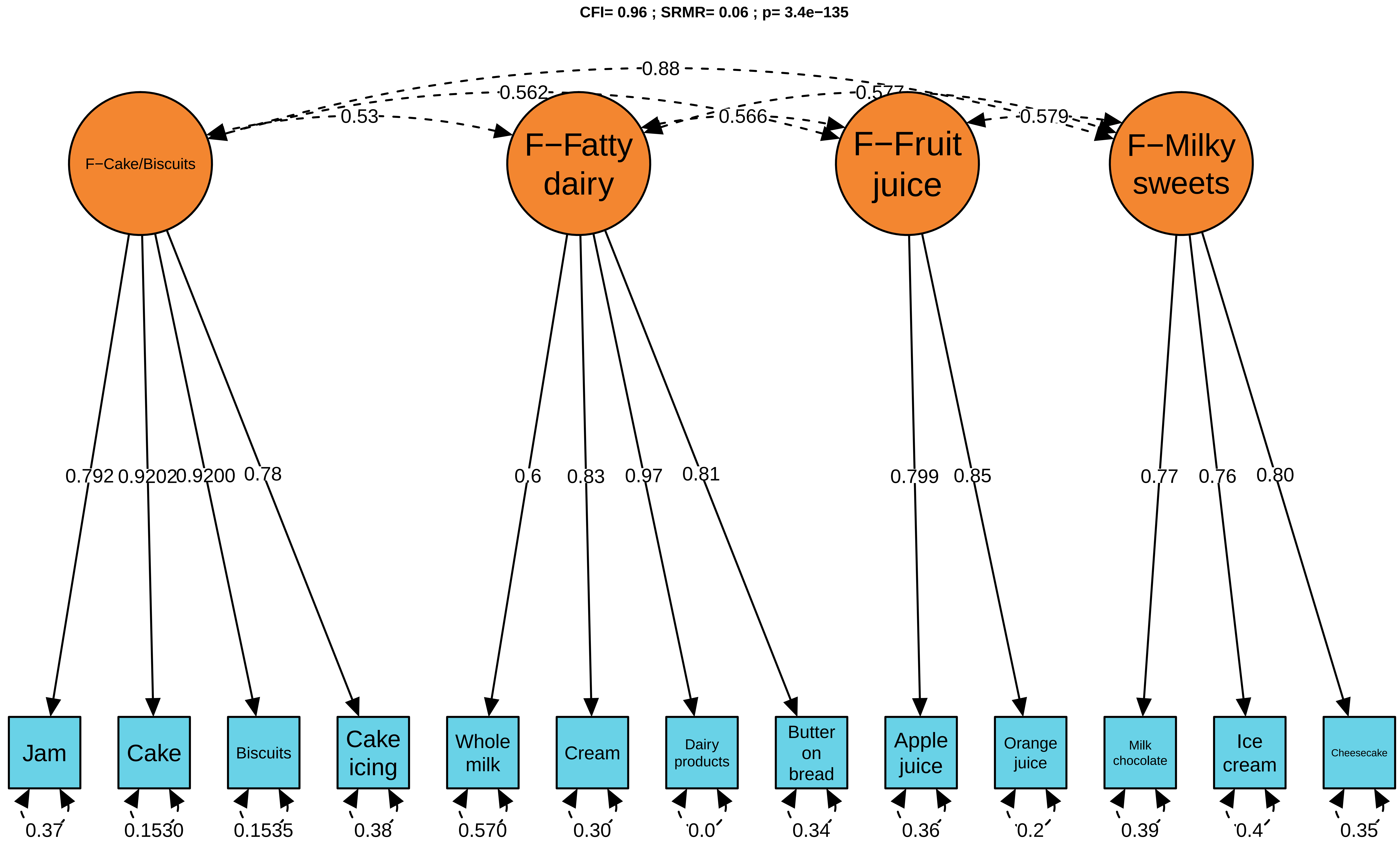
CFI= 0.96 ; SRMR= 0.06 ; p= 3.4e−135
0.88
0.562
0.53
0.577
0.566
0.579
F−Cake/Biscuits
F−Fatty dairy
F−Fruit juice
F−Milky sweets
0.792
0.9202
0.9200
0.78
0.6
0.83
0.97
0.81
0.799
0.85
0.77
0.76
0.80
Jam
Cake
Biscuits
Cake icing
Whole milk
Cream
Dairy products
Butter on bread
Apple juice
Orange juice
Milk chocolate
Ice cream
Cheesecake
0.37
0.1530
0.1535
0.38
0.570
0.30
0.0
0.34
0.36
0.2
0.39
0.4
0.35

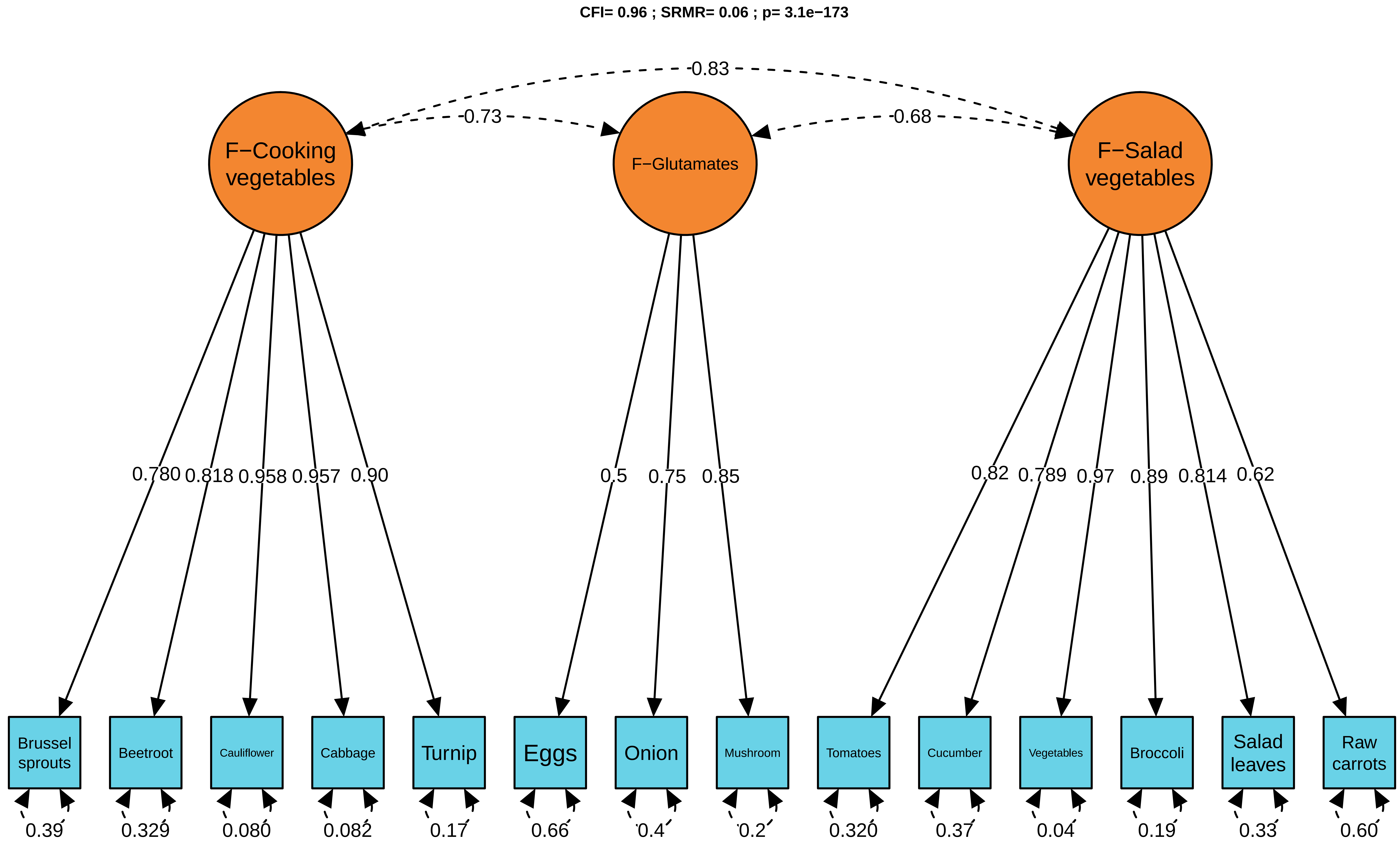
CFI= 0.96 ; SRMR= 0.06 ; p= 3.1e−173
0.83
0.73
0.68
F−Cooking vegetables
F−Glutamates
F−Salad vegetables
0.780
0.818
0.958
0.957
0.90
0.5
0.75
0.85
0.82
0.789
0.97
0.89
0.814
0.62
Brussel sprouts
Beetroot
Cauliflower
Cabbage
Turnip
Eggs
Onion
Mushroom
Tomatoes
Cucumber
Vegetables
Broccoli
Salad leaves
Raw carrots
0.39
0.329
0.080
0.082
0.17
0.66
0.4
0.2
0.320
0.37
0.04
0.19
0.33
0.60
